# Discovery of non-nucleoside inhibitors of the enterovirus D68 3D polymerase through crystallographic fragment and high-throughput biochemical screening

**DOI:** 10.64898/2026.07.09.737532

**Authors:** Iman Biswas, Qinghui Wang, Jack T. McCann, Egor P. Tchesnokov, Linh Nguyen, Manisha Saini, Jocelyn Cantero, Jezrael Lafuente Revalde, Matthias Götte, Adam R. Renslo, R. Jeffrey Neitz, Michelle R. Arkin, Eddy Arnold, Francesc Xavier Ruiz

**Affiliations:** Center for Advanced Biotechnology and Medicine, Rutgers, the State University of New Jersey, Piscataway, NJ 08854, USA; Department of Chemistry and Chemical Biology, Rutgers, the State University of New Jersey, Piscataway, NJ 08854, USA; Department of Pharmaceutical Chemistry, UCSF, CA 94158, USA; Department of Medical Microbiology and Immunology, University of Alberta, Edmonton, Alberta T6G 2E1, Canada

## Abstract

Enterovirus D68 (EV-D68) is a non-polio picornavirus that has caused increasing rates of severe respiratory illness and acute flaccid myelitis in children worldwide this century. There are no approved vaccines or antivirals for EV-D68. Thus, we conducted a crystallographic fragment screening (CFS) and a high-throughput screening (HTS) biochemical assay against the EV-D68 RNA-dependent RNA polymerase 3D (3D^pol^) to identify ligandable sites and non-nucleoside compounds that can spearhead anti-enteroviral drug discovery. The CFS, involving 650 fragments, identified 68 hit compounds (~10% hit rate) distributed across 3D^pol^, including the functionally relevant sites RNA template channel, Active site, and RNA primer channel, and the previously unknown “Thumb site II” and “Index-middle finger pocket”. Inhibition assays confirmed that compounds binding to each site can inhibit EV-D68 3D^pol^ activity. The HTS, a fluorescence-based PicoGreen biochemical assay, permitted screening 50,000 compounds of the ChemBridge Premium Library (0.77% hit rate). After a second-round dose-response screening, we identified 5-aminoindazole as a promising scaffold that inhibits EV-D68 3D^pol^, including hit-to-lead compound 727590, which displayed an IC_50_ value of 25 μM and preliminary structure-activity relationships. These hits offer amenable starting points for discovery and development of non-nucleoside inhibitors and provide opportunities for structure-based drug design against enteroviruses.

**GRAPHICAL ABSTRACT:** Created with biorender.com and PyMOL Molecular Graphics System, version 2.5.0. Schrödinger, LLC.

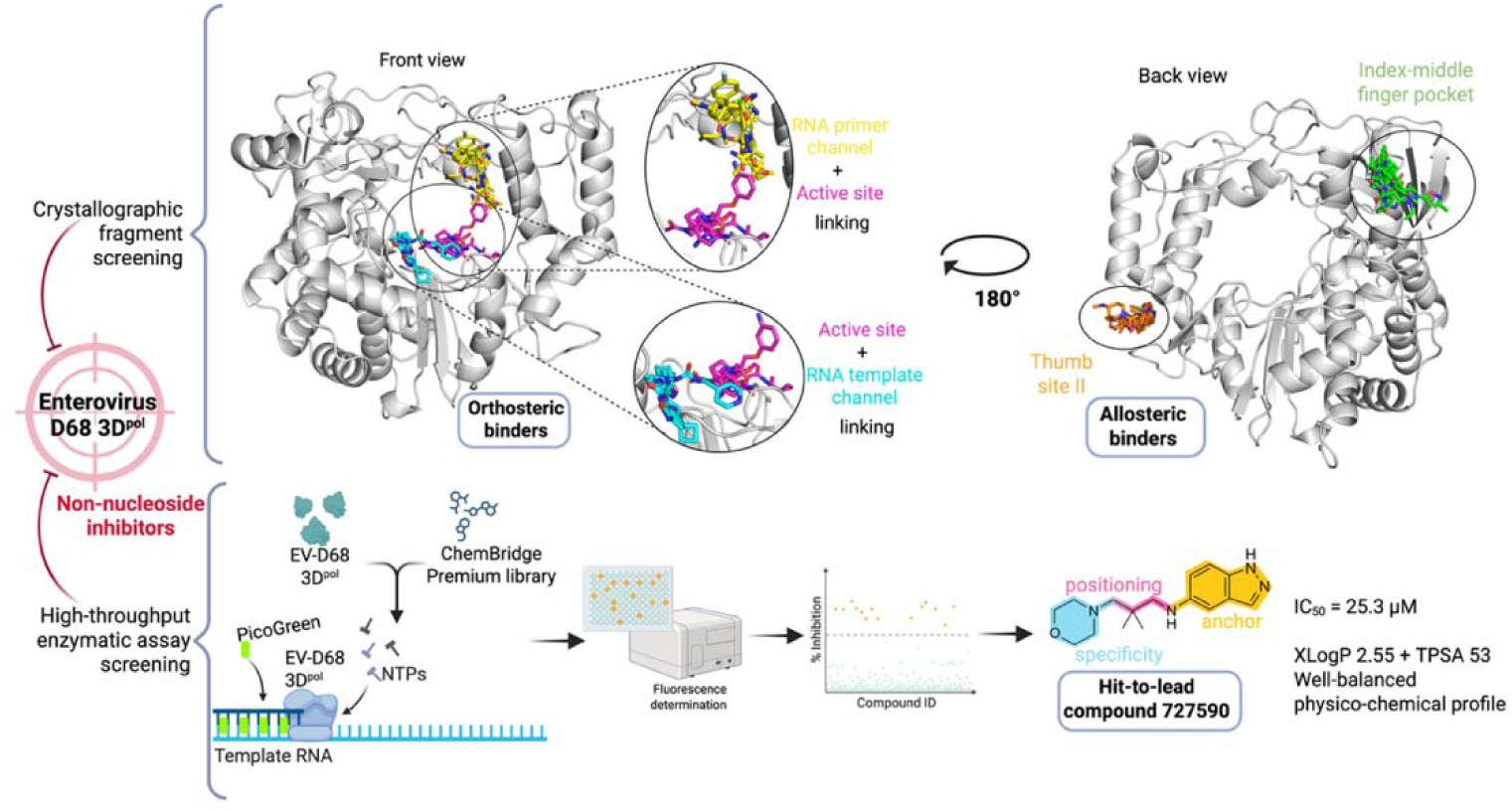

## INTRODUCTION

Enteroviruses (EVs) are small, non-enveloped positive-sense single-stranded RNA viruses, which belong to the family *Picornaviridae*. Enterovirus D68 (EV-D68) is an emerging viral pathogen belonging to the non-polio enterovirus group that has caused sporadic infections worldwide. It has primarily been detected in respiratory illnesses and is associated with occasional neurological infections (1). The pathogen can be detected in an infected person’s respiratory secretions, such as saliva and nasal mucus, hence, it can spread easily through cough and sneezes. EV-D68 was first isolated in California, USA, in 1962 and was recognized as a human pathogen responsible for various degrees of respiratory illness (2,3). Since then, many localized clusters of EV-D68 have been reported worldwide. In recent years, an outbreak occurred in Japan in 2010 with 88 cases (4). In 2012-2014, EV-D68 was detected in many children in polio-like infection clusters in California and Colorado (5). In 2014, EV-D68 caused unexpected widespread severe respiratory illness across North America with 1153 cases reported. This outbreak led to 3 deaths in Canada and 14 deaths in the USA (2).

Since 2014, EV-D68 has emerged as a quasi-biennial epidemic in the United States and Europe, with the last significant outbreak in September 2022 (6). In some cases, EV-D68 has been found to cause polio-like paralysis called acute flaccid myelitis, which is associated with the sudden onset of weakness in the arms and legs due to brainstem and grey matter lesions (7,8). EV-D68 can cause serious complications, particularly in children and those with a weakened immune system (3). Due to this increased frequency of outbreaks and lack of vaccines and antiviral drugs, it is important to develop novel therapeutics against EV-D68.

The viral RNA-dependent RNA polymerase (RdRp) 3D, also known as 3D^pol^, plays a key role in viral replication. Hence, it is highly conserved across different EV-D68 strains. 3D^pol^ initiates the synthesis of a negative-strand copy of the incoming viral genome to generate a double-stranded RNA replication intermediate. Then, the negative strand serves as a template for the synthesis of new positive-strand genome copies by 3D^pol^, which are in turn used for translation or packed into new virions (9).

RdRps catalyze RNA template-dependent formation of phosphodiester bonds between ribonucleotides in the presence of divalent metal ions (9,10). The initiation of synthesis occurs at the 3′-end of the template using a small protein primer, named VPg, and proceeds in the 5′ to 3′ direction (11). To date, a significant number of crystal structures of RdRps from the picornavirus family have been reported. These include RdRps from poliovirus, coxsackieviruses A16 and B3, enteroviruses A71 and D68, human rhinoviruses 14 and 16, and foot-and-mouth disease virus, among others (12–14).

All RdRps (including the picornaviral) adopt a similar and conserved fold, resembling a cupped right hand with fingers, palm, and thumb subdomains. The fingers can be divided into different regions: pinky, ring, middle and index (Fig. S1). Besides exhibiting overall structural similarity, RdRps display seven conserved sequence motifs, namely A-G. Motifs A-D are shared between the single-subunit RNA and DNA polymerases, motifs E-F are unique to RdRps and reverse transcriptases (RTs), while motif G is an RdRp-specific structural element that interacts with the RNA template (13). Motifs A-E are located on the palm subdomain, which contains the active site in single-subunit polymerases and represents the most conserved region, while motifs F and G are part of the fingers’ subdomain (Fig. S1). Both motifs B and F participate in nucleotide triphosphate (NTP) substrate binding, interacting with the NTP ribose and triphosphate, respectively. Picornaviral RdRps also harbor motif H in their thumb subdomain, a specific motif found in single-stranded (+) RNA and dsRNA viruses, and participate in primer strand binding and protein stability (9,15). In addition to the three subdomains and eight structural motifs, EV-D68 3D^pol^ contains three channels arrayed with positive charge: the NTP entry channel, the RNA template channel, and the RNA exit channel (15).

Crystallographic fragment screening (CFS) is a well-validated method for identifying novel compounds for the subsequent development of inhibitors (16–18). High-throughput screening (HTS) is a technique that enables rapid testing of millions of compounds for their biological activities via automated screening (19). Hit identification by CFS has the advantage of providing detailed information about the specific protein-ligand interactions and protein dynamics, as well as being very efficient at sampling the chemical space (20). Meanwhile, HTS offers the primary advantage of immense scale and functional relevance, providing immediate evidence of enzymatic inhibition or biological activity under physiological-like conditions (21).

With the aim of discovering novel non-nucleoside inhibitor scaffolds and leveraging the complementarity of both approaches, we have performed CFS and HTS campaigns targeting the EV-D68 3D^pol^. The CFS has yielded 68 hits spanning three known sites—RNA template channel, Active site, RNA primer channel—and two previously unknown sites: the “Thumb site II”, and the “Index–middle finger pocket”. The HTS, using an enzymatic inhibition PicoGreen-based assay (19), has provided 388 initial hits, including a 5-aminoindazole scaffold with hit-to-lead compound 727590, displaying an IC_50_ value of 25 μM against EV-D68 3D^pol^. The HTS enzymatic assay enabled the identification of single-site fragment hits across all five CFS binding sites. Functional validation demonstrated that fragments targeting each site inhibited enzymatic activity, confirming all newly identified sites are ligandable (except for the RNA template channel, previously validated) (15). Overall, our combined CFS and HTS campaigns provide valuable information for the design of follow-up compounds with higher binding affinity and activity inhibition of the EV-D68 3D^pol^, as well as related RdRps from other viruses from the *Picornaviridae* family.

## MATERIALS AND METHODS

### Expression and purification

The EV-D68 3D^pol^ gene was codon optimized for *E. coli* expression and synthesized by Genscript in the pET28a vector. The pET28a–EV-D68 3D^pol^ construct was transformed into *E. coli* BL21 (DE3) RIL cells and plated on LB agar with 50 μg/mL of kanamycin and 18 μg/mL chloramphenicol. A single colony was inoculated in 20 mL of Luria Broth (LB) media containing 50 μg/mL of kanamycin and 18 μg/mL chloramphenicol, followed by overnight incubation at 37 °C. This bacterial culture (A600 ~ 1) was added to one liter LB media containing 50 μg/mL of kanamycin and grown at 37 °C at 180 rpm of shaking until its optical density at 600 nm reached a value of 0.6. Protein expression was induced with 0.5 mM Isopropyl-β-D-thiogalactopyranoside (IPTG, Bio) and incubated at 16 °C for 16 hours at 180 rpm.

The cells were harvested at 4000*g* and resuspended in a lysis buffer containing, 50 mM HEPES (pH 8.0), 300 mM NaCl, 20% (v/v) glycerol, 1 mM tris(2-carboxyethyl)phosphine (TCEP) and 1 mM phenylmethanesulfonylfluoride (PMSF). The cells were lysed by sonication and then centrifuged for 45 minutes at 44,000*g* (4 °C). The supernatant was passed through nickel-nitrilotriacetic acid (Ni-NTA) beads, followed by washing the unbound protein with the same buffer supplemented with lysis buffer supplemented with 10 mM and 20 mM imidazole respectively. The protein was eluted in a buffer containing 50 mM HEPES (pH 8.0), 300 mM NaCl, 20% (v/v) glycerol, and 200 mM imidazole, diluted to reduce the NaCl concentration to approximately 100 mM prior to loading onto a HiTrap Q HP column (GE Healthcare), equilibrated with buffer containing 50 mM Tris pH 8, 100 mM NaCl, 20% glycerol, and 1 mM TCEP. 3D^pol^ was eluted using a linear salt gradient from 100 mM NaCl to 500 mM NaCl. The peak fractions were pooled, concentrated and loaded onto a Superdex 200 column (GE Healthcare) equilibrated with buffer containing 50 mM HEPES (pH 7.6), 300 mM NaCl, 3% Glycerol, 50 mM MgCl_2_, and 10 mM dithiothreitol (DTT). The eluted protein was concentrated to 12 mg/ml, aliquoted, flash frozen in liquid nitrogen, and stored at −80 °C for crystallization.

### Crystallization, structure determination, and optimization for fragment screening

Crystals were grown in a crystallization solution comprising 0.1 M Tris-HCl (pH 8.5), 16% polyethylene glycol (PEG) 3350, and 16% isopropanol by sitting-drop vapor diffusion in MRC 3 Lens Crystallization plates (SWISSCI) at 16 °C, with 0.4 nl of the aforementioned reservoir solution, and a 1:1 v/v ratio of protein to crystallization solution. Initial data collection was carried out at the SSRL BL12-2 beamline, and the structure was solved by molecular replacement using the structure of EV-D68 3D^pol^ complexed to GTP (PDB code: 5XE0, with GTP and other heteroatoms removed) as a reference model using the Phaser program in the Phenix suite (22). The solvent tolerance of the crystals was checked by incubating the crystals with 10-30% dimethyl sulfoxide (DMSO) for 1-3 hours at 20 °C and collecting the corresponding datasets also at the SSRL BL12-2 beamline.

### Fragment screening

Around 650 crystals were grown in house and carried to the XChem facility at the Diamond Light Source, UK. Due to limitation in the number of crystals, we screened 650 fragments exclusively from the DSi-Poised Library (23). Fragments were soaked into drops containing EV-D68 3D^pol^ crystals at 20 % (v/v) DMSO using acoustic dispensing with an Echo 650 liquid handler (Labcyte) (24). After incubation for 2 hours at 20 °C, the crystals were harvested using the Crystal Shifter (Oxford Lab Technologies) (25), mounted on loops and cryo-cooled in liquid nitrogen without additional cryoprotectant. Data was collected at the I04-1 beamline (Diamond Light Source, UK) at 100 K and automatically processed with Diamond Light Source’s autoprocessing pipelines using XDS (26) and either xia2 (27), autoPROC (28), or DIALS (29) with the default settings.

Analysis of fragment binding was done using XChemExplorer (30). Electron density maps were generated with DIMPLE (31), ligand restraints were calculated with ACEDRG (32) and GRADE (33), and ligand-binding events were identified using PanDDA2 (34,35). Ligands were modelled into PanDDA2-calculated event maps manually using Coot (36), and structures were refined with Refmac (37).

Coordinates and structure factors for the ground-state (PDB 14AA) and ligand-bound EV-D68 3D^pol^ structures (group deposition G_1002366) discussed here are deposited in the Protein Data Bank. Data collection and refinement statistics are summarized in Supplementary Data File 1.

### High-throughput screening assay

#### Primary screening using the fluorescence-based PicoGreen biochemical assay

EV-D68 3D^pol^ RdRp activity was determined using a fluorescence-based biochemical assay, as previously described with minor modifications (19). EV-D68 3D^pol^, expressed and purified as described above, was diluted to a final concentration of 5 μM in assay buffer containing 50 mM Tris-HCl (pH 8.0), 1 mM MgCl_2_, 0.005% Triton X-100, and PicoGreen (ThermoFisher, final concentration: 10 μl/ml). The enzyme solution was dispensed into 384-well low-volume black round-bottom microplate (Corning, Catalog No.: 4514) at2.5 μl per well using a Dragonfly liquid handler (SPT Labtech). Columns 23 and 24 were added assay buffer containing PicoGreen and remaining reagents, but not EV-D68 3D^pol^, to serve as a positive value for 100% inhibition. Next, 10 nl of each compound from the ChemBridge Premium Library (50,000 compounds) was transferred into assay plates using an Echo 650 acoustic liquid handler (Beckman Coulter Life Science), and the resulting assay plates were centrifuged at 300*g* for 1 minute, followed by a 15-minute incubation in the dark. The assay buffer containing 5 μM template (sequence: 5’-UGCGCUAGUUUUUUUUUUUUUUUUGAUCGCGU) and 200 μM dNTP mix (ThermoFisher) was then added to the plate, with a volume of 2.5 μl per well, using a Dragonfly liquid handler (SPT Labtech). Assay plates were then sealed, centrifuged at 300*g* for 1 minute, and incubated in the dark for 19 hours. Fluorescence was measured using an Envision multimode plate reader (PerkinElmer) with excitation and emission wavelengths at 485 nm and 520 nm, respectively.

#### IC_50_ determination and counterscreen using the PicoGreen biochemical assay

35 compounds identified from the primary screening were selected for IC_50_ determination and counterscreen using the same assay described in the primary screening. All compounds were evaluated in duplicates using a 10-point dose-response curve. Unlike the primary screening, in the counterscreen, an assay buffer containing PicoGreen but lacking EV-D68 3D^pol^ was treated with compounds and then incubated with the assay buffer containing 5 μM RNA duplex (sequences: 5’-ACGCGAUCAAAAAAAAAA-3’ and 5’-UUUUUUUUUUUUUUUUGA-3’) and 200 μM dNTP mix. PicoGreen binding to the RNA duplex generates a high fluorescence signal. Compounds that reduced fluorescence in the absence of 3D^pol^ were classified as false positives.

#### IC_50_ determination using a gel-based primer extension assay

To validate the inhibition results of Z56040660, compound 727590 (HTS hit), and fragments RLA-6625 to RLA-6629, gel-based *in vitro* primer extension assays were performed similarly as previously described (38). The template and primer utilized were 3’-UGCGGUACUUUUUUUUUUUUUUUU-5’ and 5’-pACGC-3’, respectively. Reactions requiring non-5′-labeled primer were supplemented with 0.1 μM [α-^32^P]-CTP for visualization. Standard reaction mixtures (15 μl) consisted of purified EV-D68 3D^pol^, assay buffer (50 mM Tris-HCl, 2.5 mM MgCl_2_, 0.005% Triton X-100, pH 8.0), 200 μM primer, 2 μM template, and NTPs. The compounds (or the DMSO control) were pre-incubated for 20 min with enzyme only, and reactions were started by the addition of template, primer and NTPs and lasted 45 min at 30 °C. Reactions were stopped with equal volumes of a formamide, 50 mM EDTA mixture. Mixtures were boiled at 95°C for 5 min, and 3 μl samples resolved by 8 M urea, 20% polyacrylamide gel electrophoresis. Radiolabeled or fluorescent signals were visualized using an Amersham Typhoon biomolecular imager and quantified in Cytiva ImageQuant TL software.

### Chemistry

Chemical reagents were purchased from commercial suppliers as indicated above and used without further purification. See the Supplementary Data section for detailed methods of the synthesized fragments.

### Figure preparation

High-resolution structural figures were prepared for publication using PyMOL (The PyMOL Molecular Graphics System, Version 2.5, Schrödinger, LLC) and BioRender (biorender.com).

## RESULTS

### Multiple Binding Sites on EV-D68 3D^pol^ identified by crystallographic fragment screening

We soaked 650 fragments into EV-D68 3D^pol^ apo crystals, with 498 datasets successfully collected, 98% of which diffracted to at least 2.0 Å, with an average resolution of 1.5 Å overall. We have identified 68 fragment hits accounting for 81 binding events, with 55 single binders, 12 fragments bound at two sites, and 1 fragment bound at three sites (Fig. 1A). Some of the hits are in the RNA binding cleft of the EV-D68 3D^pol^ while some are on the surface of the protein (Fig. 1B). PDB IDs, names, ligand descriptors, and data collection and refinement statistics are available in Table S1.

**Figure 1:**
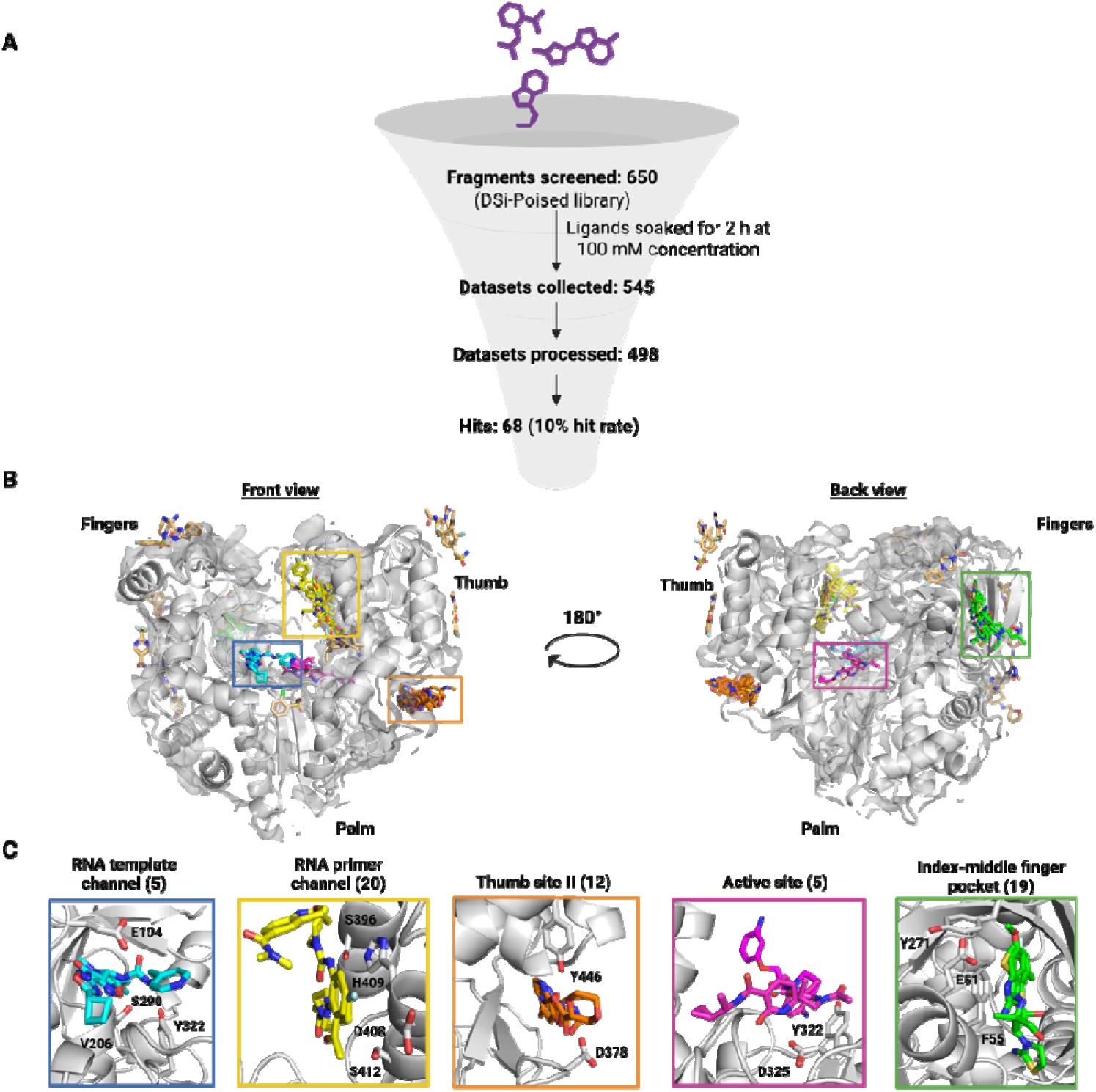
Crystallographic fragment screening campaign of EV-D68 3D^pol^ reveals orthosteric and allosteric binding sites. A) Overview of the crystallographic fragment screening campaign presented in this study. B) Front and back view of the EV-D68 3D^pol^ structure with all the fragment hits. Fragments bound to the five sites identified boxed in different colors: RNA template channel (blue); RNA primer channel (yellow); Thumb site II (orange); Active site (magenta); Index-middle finger pocket (green). The remaining unassigned fragments are displayed in light brown. C) Closer view of the binding sites boxed in (B), with the name of the site and the number of hits in parenthesis. All the fragment hits are depicted a sticks, and the protein backbone is represented as grey cartoon and surface. Created with biorender.com and the PyMOL Molecular Graphics System, version 2.5.0. Schrödinger, LLC.

Based on their location within the RNA binding cleft and on the literature on RdRps from the *Picornaviridae* family, we have identified three sites with potential of driving the development of non-nucleoside small molecule inhibitors: 1) the RNA template channel, 2) the Active site and, 3) the RNA primer channel. Both the RNA template channel and the Active site bind five hits, while the RNA primer channel is a hotspot interacting with twenty fragments (Fig. 1B-C).

In addition, two other hot spot binding sites outside the RNA binding cleft have been identified: 1) Thumb site II, with twelve binders, and 2) Index-middle finger pocket, with nineteen binders (Fig. 1C). Fig. S2 displays all hits grouped in the previous five sites. Additionally, we have identified 7 fragment hits dispersed at other regions of the EV-D68 3D^pol^.

### Fragment hits in the RNA template channel

We have identified five fragment hits in the RNA template channel (Fig. 2A-E). Upon superimposition of the fragment bound structures of EV-D68 3D^pol^ with the structure of the EV-A71 3D^pol^ elongation complex (PDB 6KWR), we observe that the five fragment hits are overlapping with a portion of the RNA template strand (Fig. 2F), hence we identified this binding site as the RNA template channel.

**Figure 2:**
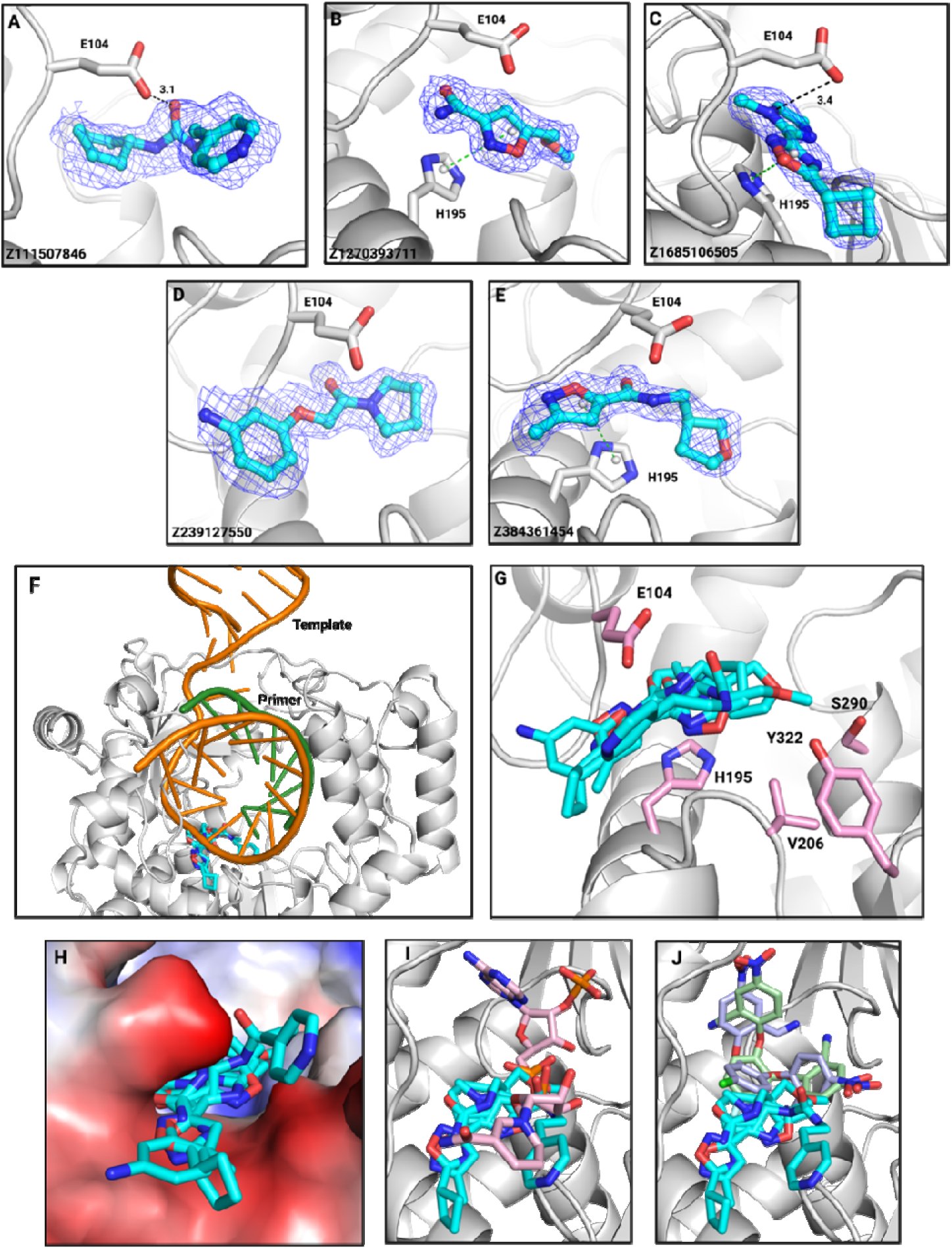
Fragment hits in the RNA template channel partially overlap with known EV 3D^pol^ inhibitors. (A-E): Distinct fragments bound to the RNA template channel. Hydrogen bonds and polar interactions are displayed in black dashed lines, while stacking interactions in green dashed lines. F) Fragment hits (cyan) overlapping with the RNA template (orange, RNA primer in green), by superposition with the EV-A71 3D^pol^ elongation complex (6KWR). G) Overview of the RNA template channel with fragment hits (cyan) and side chains of the key residues (pink) in the pocket. H) Electrostatic charge distribution at the RNA template channel site. EV-D68 3D^pol^ structure with fragment hits (cyan) at the RNA template channel superimposed to: I) NADPH bound (pink) structure of EV-D68 3D^pol^, J) GPC-N143 (green) and GPC-N114 (pale blue) bound structure of CVB3 3D^pol^. Created with biorender.com and the PyMOL Molecular Graphics System, version 2.5.0. Schrödinger, LLC.

There are previous reports of small molecule inhibitors binding to the RNA template channel of picornaviral RdRps. The key biosynthetic cofactor NADPH (Table S2) was identified as EV-D68 3D^pol^ inhibitor, with an IC_50_ value of 232.9 µM in recombinant enzyme inhibition assays and was found to occupy the RNA template channel of the enzyme in the crystal structure of the complex (15). There are also cocrystal structures with two other small molecule inhibitors, GPC-N114 and GPC-N143 (Table S2), bound to the RNA template channel of coxsackievirus B3 RdRp, with GPC-N114 exhibiting antiviral activity against a panel of enteroviruses ranging 0.1 to 2.0 µM EC_50_ values, including EV-D68 with an EC_50_ of 1.44 µM (39).

This RNA template channel portion surrounding the fragment hits is lined with amino acid residues, G102, L103, E104, V206, G207, S290 and Y322 (Fig. 2G, H), making it mostly hydrophobic (except for the entry part, Fig. 2G). These residues are highly conserved in picornaviruses (Fig. S4). Nevertheless, we observe hydrogen bonding or polar interactions between the side chain of E104 residue and the fragment hits, and π-π stacking interactions between the side chain of H195 and the aromatic ring of some of the fragment hits (Fig. 2A-E).

This portion of the site partly overlaps with the binding site of NADPH in its EV-D68 3D^pol^ cocrystal structure (Fig. 2I). Upon superimposition of the fragment-bound structures of EV-D68 3D^pol^ to the small molecule inhibitor-bound structures of Coxsackievirus B3 RdRp, we observed that there is a partial overlap of the binding pockets. However, the fragment hits cover the entry portion of the tunnel, while the small molecule hits span across the inner part of it (Fig. 2J).

### Fragment hits in the Active site

We have identified five fragment hits at the Active site (Fig. 3A-E). The active site regions in RdRps from the family *Picornaviridae* are structurally conserved across different viruses despite slight variations in their sequences (Fig. S2). The Active site sits on the palm domain, and is built on a three-stranded β-sheet composed of one strand from motif A and two strands from motif C that are packed against a long α-helix from motif B. The GDD motif (G323, D324 and D325, in the β-turn of motif C, Fig. S2) in EV-D68 3D^pol^ is the catalytic center for RdRps.

**Figure 3:**
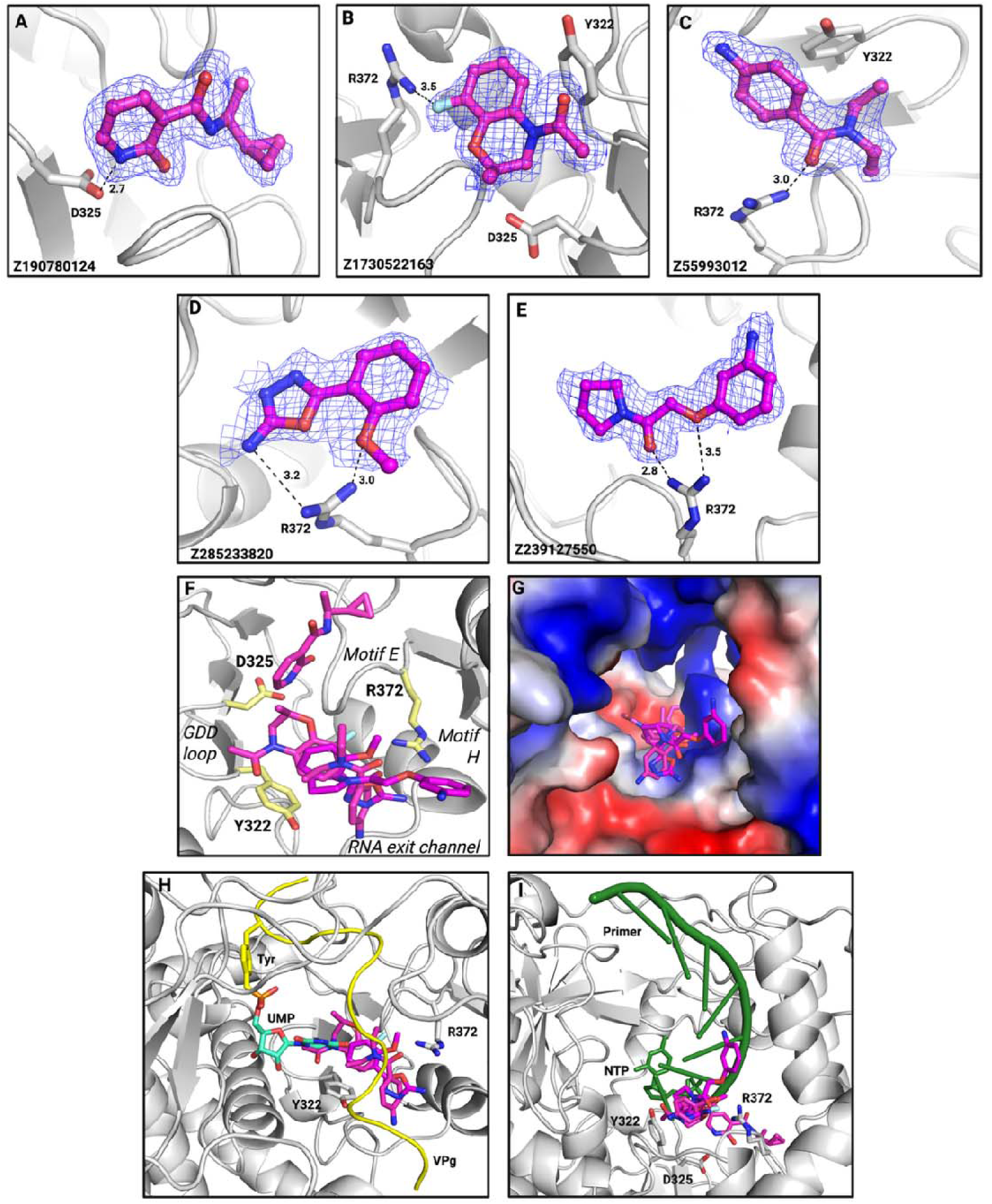
Fragment hits bind the Active site, including direct interaction with the catalytic residue D325. (A-E): Distinct fragments bound to the Active site. F) Overview of the active site (top view) with fragment hits (magenta) and side chain of the key residues (yellow) in the pocket. The front part (as in Fig. 1B left side) is formed by the palm loop located between α-helices α7 and α8 (Fig. S2), motif E (palm) is in the back, while the GDD loop of motif C and motif H (thumb) flank each side. G) Electrostatic charge distribution at the active site (front view). EV-D68 3D^pol^ structure with fragment hits (magenta) at the active site. H) The same fragment hits at the active site superimposed onto the structure of the FMDV RdRp complexed to VPg (2F8E) displaying the product of the VPg uridylylation, the VPg-UMP and I) superimposed onto the structure of the EV-A71 3D^pol^ elongation complex (6KWR) displaying the primer overlapping with the active site. Created with biorender.com and the PyMOL Molecular Graphics System, version 2.5.0. Schrödinger, LLC.

A key structural feature of the picornaviral active site is that it is wide to facilitate the protein-mediated initiation of replication using VPg (40). The RNA replication elongation process in picornaviruses has been extensively studied using poliovirus 3D^pol^ as model polymerase (41). Overall, 3D^pol^ forms a very stable and highly processive elongation complex with a well-defined catalytic cycle. Conversely, structural studies of initiation have been elusive. Additionally, the 3D^pol^ structures in the absence or presence of RNA are almost identical, not offering any insight into a potential conformational “switch” between initiation and elongation (14).

The binding site of the fragment hits is lined by residues G207, C208, Y322, D325, D354, T368, L370, K371, R372 (Fig. 3F), overall imparting both positive and negative charges to the pocket (Fig. 3G). All residues lining this site are conserved in picornaviruses (Fig. S4). Z190780124 sits on top of motif E (Fig. S1) and interacts with the side chain of the catalytic site D325 of the GDD motif by a strong hydrogen bonding interaction (donor-acceptor distance 2.7 Å, Fig. 3A). Inhibitors binding within the GDD motif might interfere with the catalysis step and block the RNA replication process. The other fragment hits at this site are located closer to the front part of the palm subdomain, near the RNA exit channel, delimitated by helices α7 and α8 (Fig. 3F). They form hydrogen-bonding interactions with the side chain of R372 and strong van der Waals interactions with the side chain of Y322 (Fig. 3B-E).

Comparison with 3D^pol^ structures in complex with RNA or VPg helps to rationalize how inhibitors binding to the Active site may hinder RNA replication. Upon superimposition of the fragment bound structures of EV-D68 3D^pol^ with the structure of the related foot-and-mouth disease virus 3D^pol^ in complex with uridylylated VPg protein (PDB 2F8E) (42), we observed that our fragment binding site shows significant overlap with the interaction site of the uridylylated VPg protein (Fig. 3H). In addition, on superimposing the fragment-bound structures of EV-D68 3D^pol^ with the structure of the EV-A71 3D^pol^ elongation complex (PDB 6KWR), some of the fragment hits also overlap with the terminal portion of the primer strand and the NTP binding site (Fig. 3I).

### The RNA primer channel is a fragment binding hotspot

We have identified twenty fragment hits at the RNA primer channel, the site with most bound hits and hence a hotspot (see representative binders in Fig. 4A-J). We have dubbed this site as RNA primer channel because it overlaps with several nucleotides of the primer strand (Fig. 4H). In fact, it is a large and relatively enclosed subpocket within the RNA binding groove of 3D^pol^. It is delimited in one side by residues in loops from the fingers’ subdomain regions index and ring (L21, residues 155 to 158, Fig. S1) and the thumb α-helices α14 and α15, parallel to each other and connected by a turn (residues 386 to 416, Fig. S1). Helix α15 forms part of the motif H of the thumb subdomain in single stranded (+) RNA and dsRNA viruses and is engaged in extensive interactions with the primer strand (Fig. 4I). The residues of this binding pocket are well conserved in picornaviruses (Fig. S4), which indicates the possibility of developing broad-spectrum inhibitors targeting this site.

**Figure 4:**
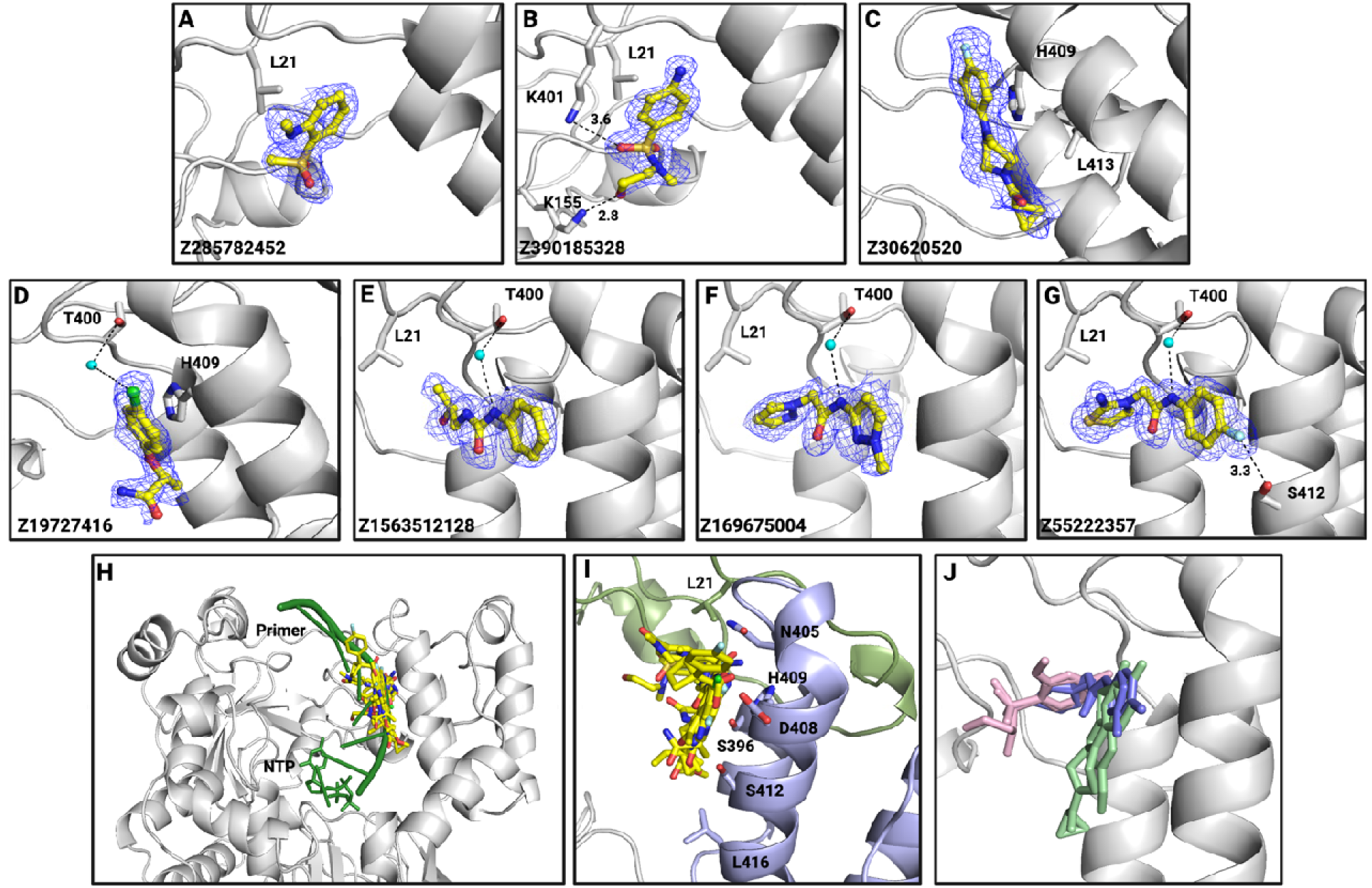
The RNA primer channel is a fragment binding hotspot. Fragments bound to the RNA primer channel, (A-B): near the fingers, (C-E): near the thumb, (F-H): bridging fingers and thumb. I) Structure of the EV-A71 3D^pol^ elongation complex (6KWR) displaying the primer overlapping with the RNA primer channel. J) Overview of the RNA primer channel showing the thumb (violet) and fingers (green) and fragment hits (yellow) with the side chain of the key residues displayed. K) Representatives of the three types of fragment binders: near fingers (pink), near thumb (green) and bridging fingers and thumb (violet). Created with biorender.com and the PyMOL Molecular Graphics System, version 2.5.0. Schrödinger, LLC.

Based on the proximity to the conforming subdomains, we can divide the fragment hits of this site in three binding types (Fig. 4J): I) near the fingers (Fig. 4A-B), II) near the thumb (Fig. 4C-D), or III) bridging fingers and thumb (Fig. 4E-G). The fragment hits type I present as main interactions a hydrogen bond with the side chain amino of K401 and a hydrophobic interaction with L21, while are surrounded on one side by the fingers’ residues 155 to 158 and on the other by thumb’s residues 396 to 401 (Fig. 4A-B, J). The fragment hits type II mainly share as common interactions hydrophobic contacts with H409 and L413, while nestled against the thumb α-helices α14 and α15 (Fig. 4C-D, J).

The type III fragment hits share as common contacts a water-meditated hydrogen bond with the oxygen atom of T400’s side chain, as well as a hydrophobic contact with L21 (Fig. 4E-G, J). The water-mediated contact is also observed with fragment Z1201620232 of type II binders (Fig. 4D). Type III hits are surrounded by residues of both the fingers and thumb subdomains (Fig. 4J).

We have observed that the fragment hits of the RNA primer channel of EV-D68 3D^pol^ partly overlap with the binding site of several small molecule inhibitors of human norovirus RdRp (of the related *Caliciviridae* family) cocrystals (Fig. S3). These include RdRp inhibitor suramin-related compounds (43) and two naphthalene-sulfonate derivatives: NAF2 and PPNDS (Table S2). PPNDS displays an IC_50_ value of 0.45 µM against recombinant human norovirus RdRp (44).

### Two sites outside of the nucleic acid binding grooves are hotspots: “Thumb site II” and “Index-middle finger pocket”

The three previous binding sites are validated targets for inhibition, as all are located within the nucleic acid binding groove, very conserved due to its importance for catalysis, and present overlap with small molecule inhibitors of EV-D68 3D^pol^ or related RdRps. Additionally, we have identified two binding sites outside of the nucleic acid binding groove which could be of potential significance, as both are hotspots.

#### Thumb site II

There are 12 fragment hits bound at the Thumb site II (Fig. 5A-F). This site is located at the base of the thumb domain, in the C-terminal region of 3D^pol^, and overlaps with small molecule inhibitors and fragments found in the “Thumb site II” of RdRps from hepatitis C virus (HCV) (45) and dengue virus (46). For HCV RdRp, Thumb site II binders include inhibitors such as filibuvir (45), lomibuvir (45), and radalbuvir (45) (Table S2 and Fig. S6), that have reached phase II clinical trials. The Thumb site II seems to regulate the motion of the β-loop in the HCV RdRp-RNA complex (Fig. 5H) required for the transition from initiation to elongation (47,48).

**Figure 5:**
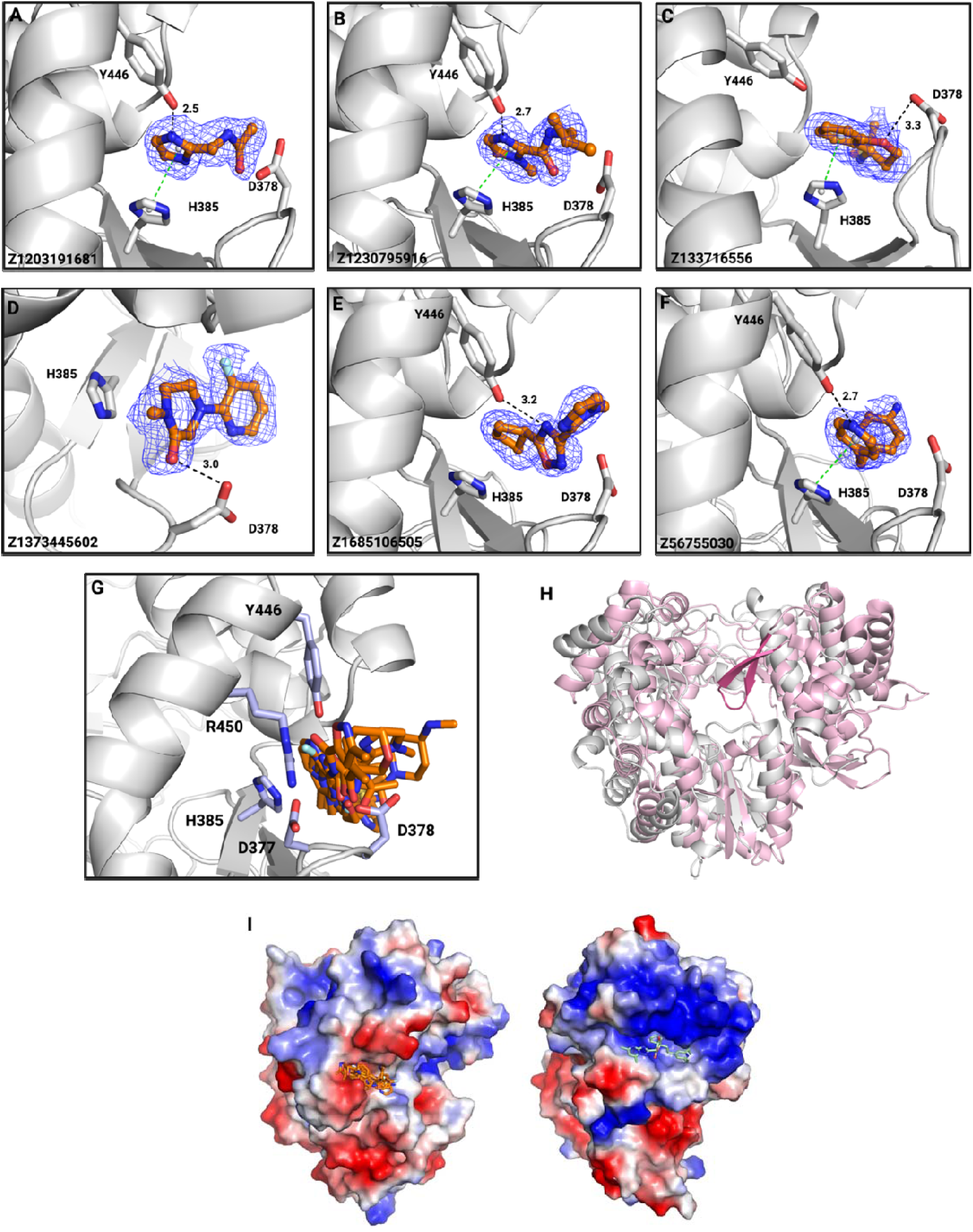
The novel site “Thumb site II” is a hotspot located in the same region but with different architecture than the analogous flaviviral and hepaciviral RdRp’s Thumb site II. (A-F): Distinct fragments bound to the Thumb site II. G) Overview of the Thumb site II with fragment hits (orange) and side chain of the key residues (pale blue) in the pocket. H) Superposition of EV-D68 3Dpol (grey) with HCV RdRp (light pink), with the priming loop of HCV RdRp highlighted in dark pink. I) Electrostatic charge distribution on the surface of EV-D68 3Dpol (left) bound to thumb site II fragment hits (orange), and HCV RdRp (right) bound to filibuvir (pale green). Created with biorender.com and the PyMOL Molecular Graphics System, version 2.5.0. Schrödinger, LLC.

Additionally, in the dengue virus RdRp, closely related to the HCV counterpart, the Thumb site II is a binding site for fragments and small molecules, as well as for the RNA stem-loop structure from the 5’-UTR region of the viral genome. Nevertheless, in picornaviruses, the priming loop is absent (Fig. 5H), and the initiation of RNA replication is primed by the VPg protein. It is also important to point out that while in HCV and dengue RdRps the thumb site II is positively charged, in the case of EVD68 3D^pol^ the site is negatively charged (Fig. 5I). In fact, the C-terminal region of the flaviviral and hepaciviral RdRps corresponds to an extension that is absent in the smaller picornaviral and noroviral RdRps. Therefore, while there is overlap in the location of the site, the functional role seems clearly divergent.

This site is moderately conserved in picornaviruses (Fig. S2). The residues lining this pocket are, H385, D377, D378, W418, H419, G421, Y446 and R450 (Fig. 5G). The most common interactions that we observe in this pocket are, the π-π stacking interaction mediated by the side chain of H385 with the aromatic ring of most of the fragment hits and hydrogen-bonding interaction between the side chain of Y446 with some of the fragment hits (Fig. 5A-F). Additionally, there are hydrophobic interactions between the fragment hits and several residues lining the pocket.

#### Index-middle finger pocket

The second binding hotspot outside of the nucleic acid binding groove is located at the junction of the middle finger and the index finger domain, which we have called “Index-middle finger pocket”. We have identified 19 fragment hits at this binding pocket (Fig. 6A-F).

**Figure 6:**
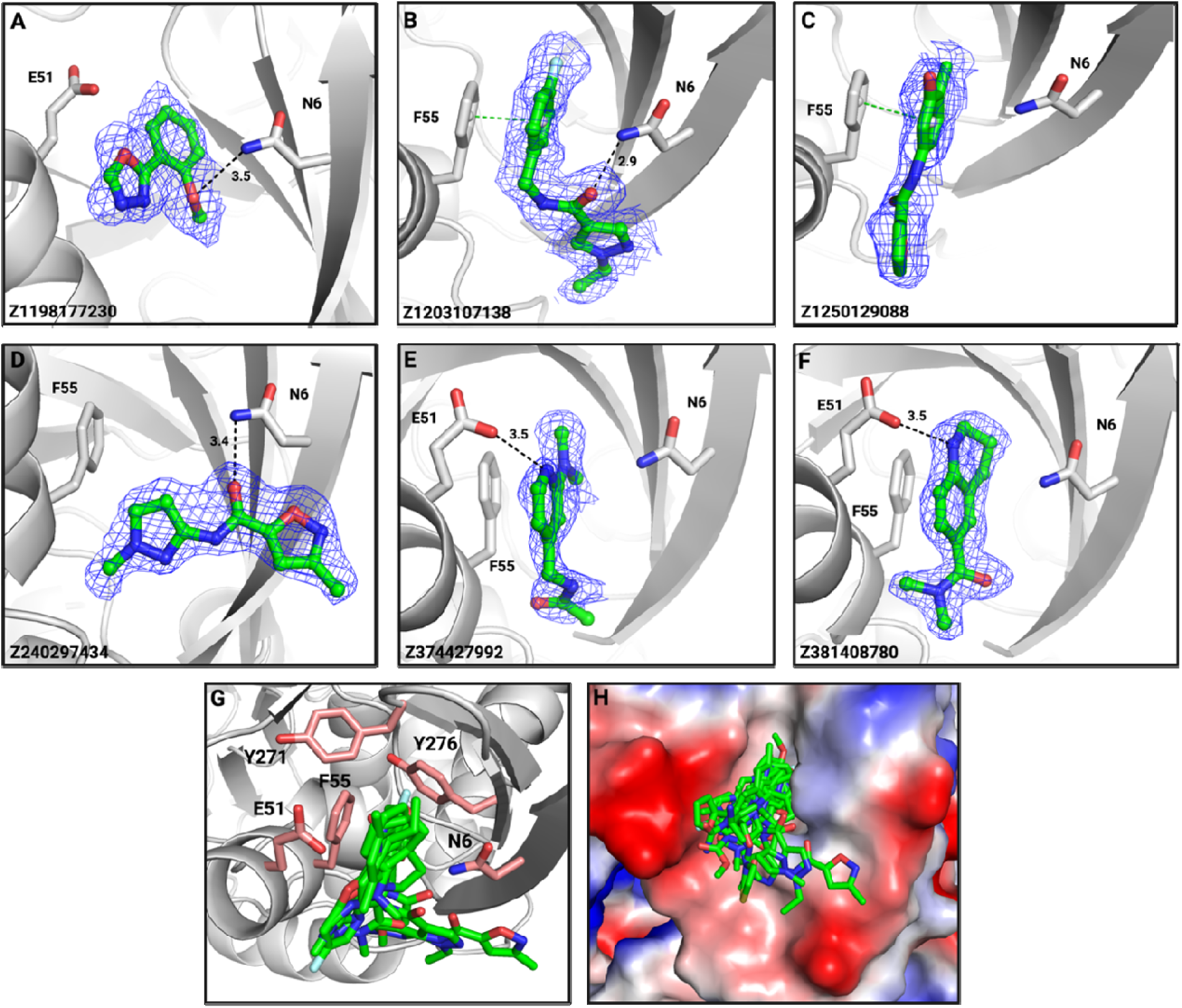
The novel “Index-middle finger pocket” is a hotspot located in the back side of EVD68 3D^pol^. **(A-F):** Distinct fragments bound to the Index-middle finger pocket. G) Overview of the Index-middle finger pocket with fragment hits (green) and side chain of the key residues (pink) in the pocket. H) Electrostatic charge distribution at the Index-middle finger pocket. Created with biorender.com and the PyMOL Molecular Graphics System, version 2.5.0. Schrödinger, LLC.

The residues in this pocket are, I3, N6, E51, F55, R274, Y271 and Y276 (Fig. 6G), overall imparting a balance of hydrophobicity, and positive and negative charges to the pocket (Fig. 6H). This pocket is moderately conserved in picornaviruses (Fig. S2). The predominant interaction at this site are the hydrophobic interactions of the fragment hits with the nearby residues. Other interactions at this site include hydrogen-bonding interactions mediated by N6 and E51 with some of the fragment hits and π-π stacking interactions mediated by the side chain of F55 with the aromatic ring of some of the fragment hits (Fig. 6A-F). Two of the interacting residues, I3 and N6 are located on the long loop of the finger domain that bridges the finger and thumb domains together and is important for the structural integrity of the enzyme (49).

### High-throughput screening assay identifies a 5-aminoindazole scaffold inhibiting EV-D68 3D^pol^

We developed and optimized conditions for an HTS PicoGreen-based EV-D68 3D^pol^ inhibition assay, using reference (19) as a starting point. Indeed, PicoGreen binds dsRNA and emits a fluorescent signal proportional to its concentration, enabling the identification of RdRp inhibitors, particularly in high-throughput screening (Fig. 7A).

**Figure 7:**
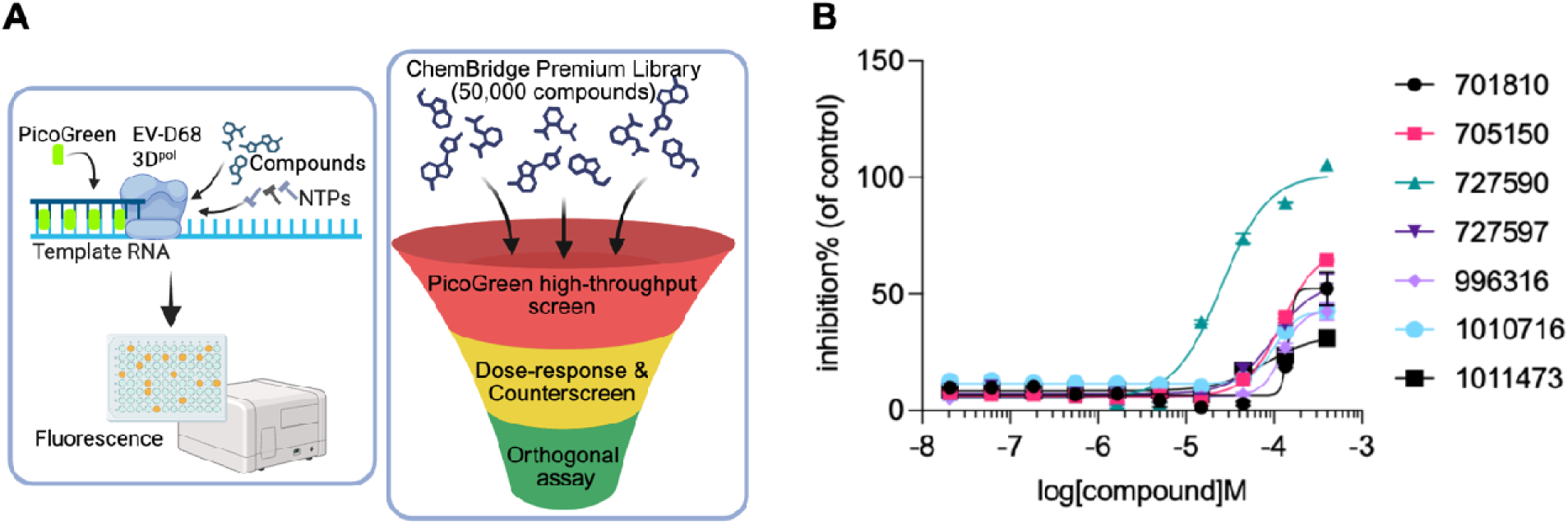
Overview of the high-throughput screening (HTS) campaign screening against EV-D68 3D^pol^. (A) Schematic of the PicoGreen-based HTS assay, and HTS pipeline. (B) Dose–response validation of the selected seven hits. Created with biorender.com.

An important condition for an HTS assay is the temperature. While 37°C is essential for maintaining cell viability in cell-based assays, room temperature (often 20–25°C) is generally superior for biochemical assays, liquid handling, and overall assay robustness (50). The optimal temperature for RdRp activity usually ranges from 30 to 33 °C (51). To assess whether an HTS assay for EV-D68 3D^pol^ can be established at room temperature, we first measured fluorescence over 30 hours by incubating 200 μM primer and 10 μM template with varying concentrations of EV-D68 3D^pol^ with a total volume of 5 μl per well.

The result indicated that the rate of fluorescence change is generally proportional to EV-D68 3D^pol^ concentration. Although fluorescence changes in a linear fashion when a low concentration of EV-D68 3D^pol^ is applied, enough EV-D68 3D^pol^ is required to produce a significant fluorescence increase. Since 2.5 μM EV-D68 3D^pol^ showed both a significant change in fluorescence intensity and linear kinetics, we used this concentration and further optimized the template and primer concentrations to minimize material use and preserve assay performance. The latter is reflected in the Z’ value: a dimensionless parameter that measures the separation between the positive control (maximum signal) and negative control (minimum signal) relative to the data variability (52). Gratifyingly, we found the combination of 2.5 μM EV-D68 3D^pol^ and 2.5 μM template demonstrated a Z’ of sufficient robustness for HTS, with a value of 0.47. Additionally, the signal was not significantly affected by DMSO, even at 4%. The chosen conditions represent a *de novo* inhibition assay. This is not representative of the *in vitro* scenario, but it was chosen for its cost-effectiveness and high-throughput scalability and has been previously described for the related poliovirus 3D^pol^ (53). Additionally, it allows targeting the conformation of EV-D68 3D^pol^ during the initiation phase of the viral cycle, which might be more sensitive to inhibition than the more processive elongation phase (14).

We then screened a total of 50,000 compounds of the ChemBridge Premium library (providing high structural diversity and favorable physicochemical properties, enabling both “drug-like” and “lead-like” hits) at a single dose of 20 µM (Fig. 7A). Overall, the screening shows high quality, with Z’ values across all tested plates ranging from 0.5 to 0.8. 388 compounds were identified as initial hits (a “hit” was defined as a data point that lies more than three standard deviations away from the mean), yielding a hit rate of 0.77% (Fig. S10A). We then clustered the initial hits based on their chemical structures and calculated their common drug-like properties. Consequently, 35 out of 388 compounds were selected for a dose-response assay and counterscreen. The counterscreen (performed with all the primary assay components except EV-D68 3D^pol^) results support the notion that none of the 35 compounds interferes with the interaction between PicoGreen and the RNA duplex (Fig. S10B).

Amongst the 35 selected hits from the primary screening, we confirmed that 7 of them dose-dependently inhibit RdRp activity in the PicoGreen assay (Fig. 7B and Table 1). Notably, compound 727590, with a 5-aminoindazole core, displayed the most potent EV-D68 3D^pol^ RdRp inhibitory activity, with an IC_50_ of 25.3 µM (Table 1). Interestingly, two other compounds, 705150, with the same 5-aminoindazole core and 727597 with a similar core structure, also showed RdRp inhibition (Table 1), although to a much weaker extent (IC_50_ values of 98 and 122 µM, respectively). Taken together, 5-aminoindazole may represent a promising scaffold to inhibit EV-D68 3D^pol^.

**Table 1.**
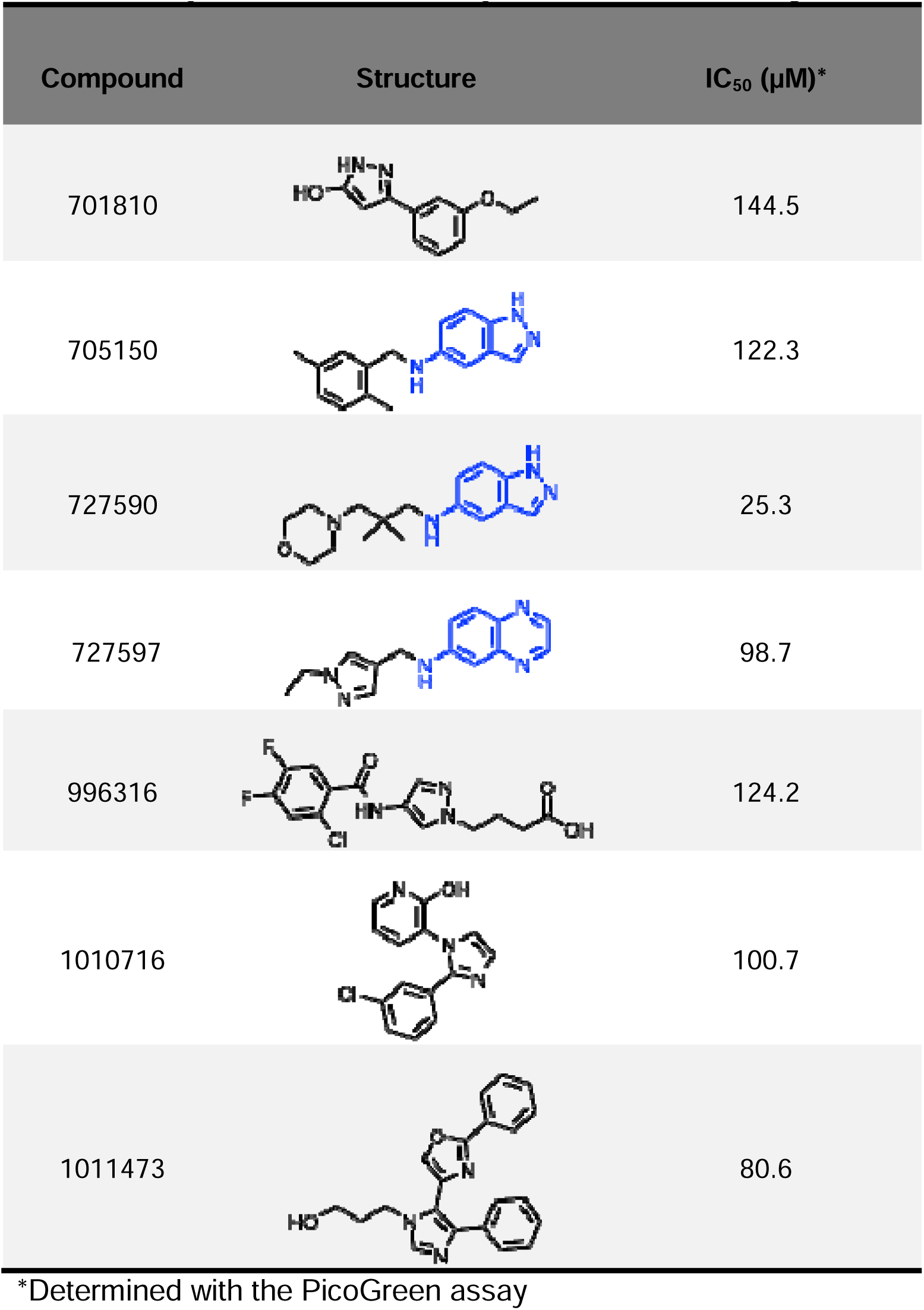
Enzyme inhibition activity of the HTS screening hits.

| Compound | Structure | IC<sub>50</sub> (μM)* |
| --- | --- | --- |
| 701810 |  | 144.5 |
| 705150 |  | 122.3 |
| 727590 |  | 25.3 |
| 727597 |  | 98.7 |
| 996316 |  | 124.2 |
| 1010716 |  | 100.7 |
| 1011473 |  | 80.6 |

\*Determined with the PicoGreen assay

To confirm this, we cherry-picked 13 additional compounds with either a 5-aminoindazole or a 6-aminoindazole core from the ChemBridge Premium and ChemBridge Gallo libraries and measured their IC_50_ values. The results are shown in Table S3. Compared with the 5-aminoindazole counterparts (46706 and 47508), the 6-aminoindazole compounds 46388 and 47389 displayed weaker EV-D68 3D^pol^ RdRp inhibitory activity, pointing to the 5-aminoindazole scaffold being preferred to the 6-aminoindazole.

### Selected single-site fragment inhibition assays show that all fragment-bound sites are inhibitory

To probe the inhibitory potential of the CFS-discovered fragment binding sites, we selected single-site fragment hits for the enzyme inhibition assay developed for HTS, described in the previous section. Hits at the Active site, Thumb site II, and Index-middle finger pocket displayed EV-D68 3D^pol^ inhibition at the single dose of 5 mM (Fig. 8A and C). From these, Z26333448 (Thumb site II) and Z56040660 (RNA primer channel) were assayed in dose-response assays (Fig. 8B) and counterscreened (Fig. S11). While Z26333448 was confirmed as Thumb site II inhibitor, with an IC_50_ value of 2.8 mM, Z56040660 was found to interfere with the signal and cannot be confirmed an inhibitor. Thus, in order to validate Z56040660 as well as the rest of PicoGreen inhibition hits, we performed a gel-based radioactivity primer extension assay (see Methods section for details). The results are summarized in Table 2 and in Fig. S12. They validate Z56040660 as well as the rest of PicoGreen inhibition hits, with a good correlation between both methods in terms of inhibition values/percentage.

**Figure 8:**
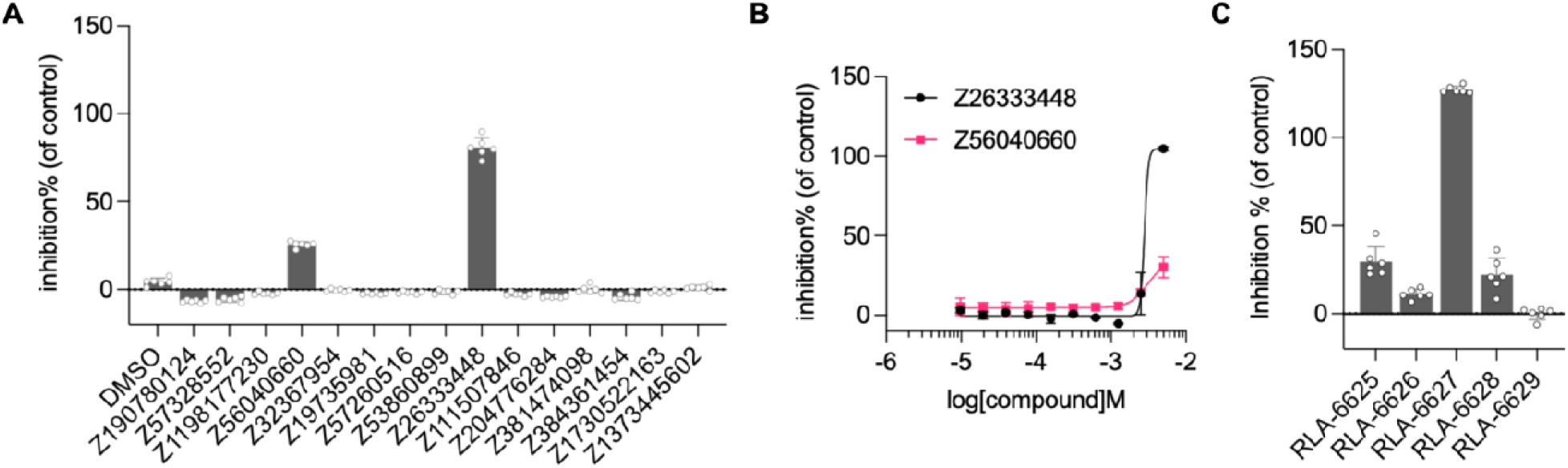
Several single-site fragment hits inhibit EV-D68 3D^pol^. (A) Selected single-site fragment hits identified from the crystallographic fragment screening campaign were assayed against EV-D68 3D^pol^ using the PicoGreen assay at a single dose of 5 mM. (B) Dose–response validation of Z26333448 (Thumb site II) and Z56040660 (RNA primer channel). (C) Selected single-site fragment hits (or related) were re-synthesized, and assayed against EV-D68 3D^pol^ using the PicoGreen assay at a single dose of 5 mM: RLA-6625: Index–middle finger pocket; RLA-6626 and RLA-6627: R & S isomers of Z190780124: Active site; RLA-6628: derivative of the previous; RLA-6629: Active site. Created with biorender.com.

**Table 2.**
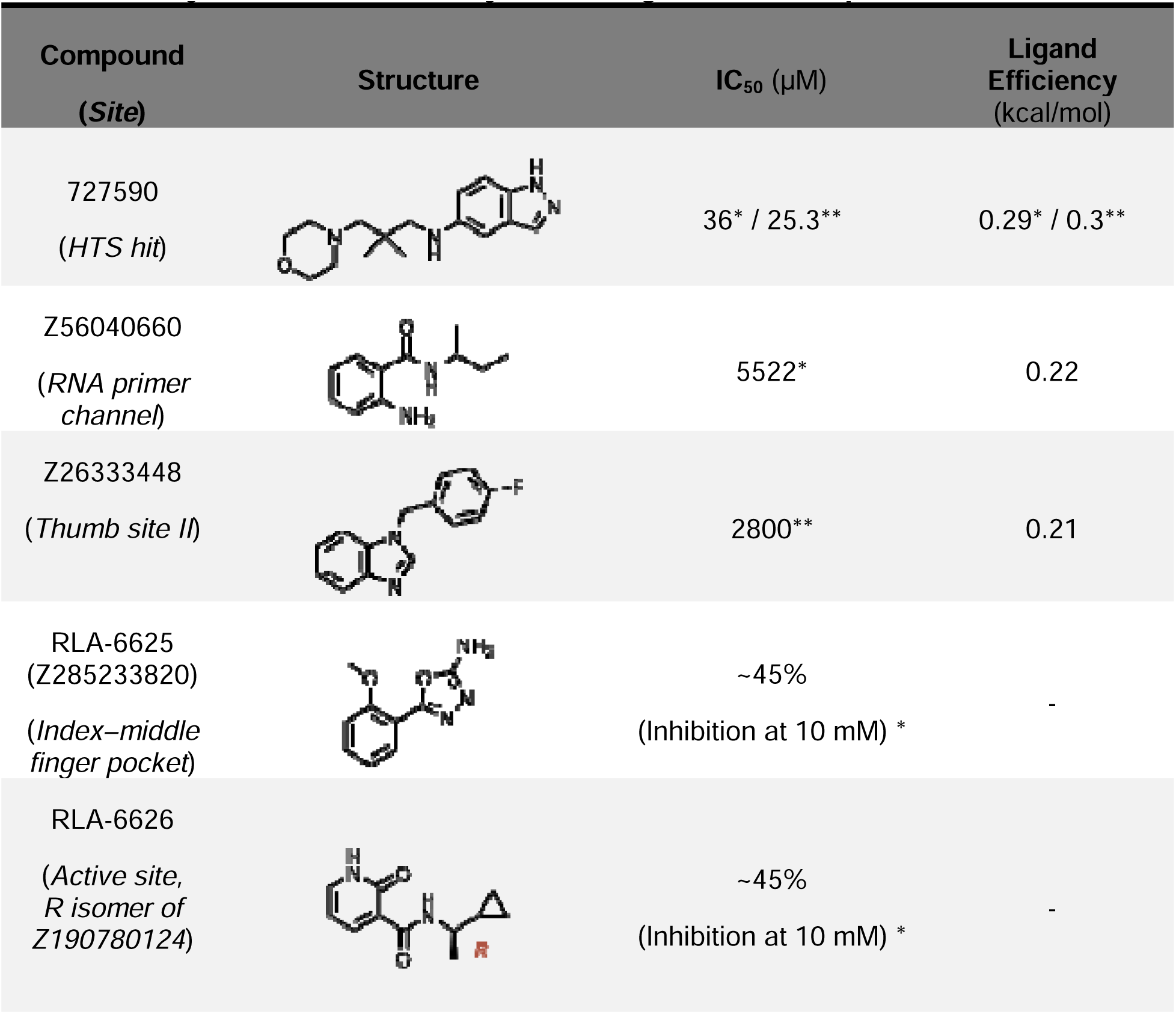
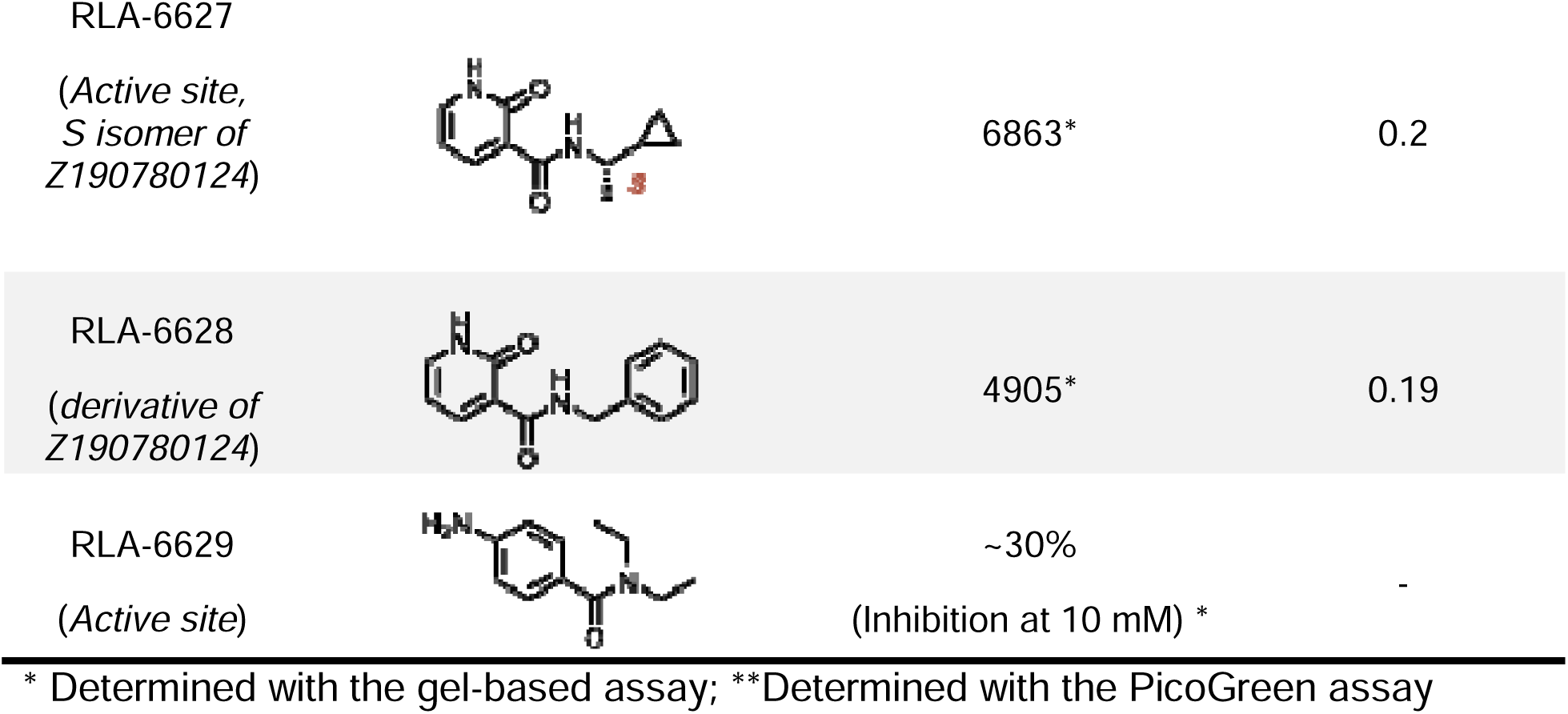
Enzyme inhibition activity of the fragment hit compounds and 727590.

| Compound<br>(Site) | Structure | $IC_{50}$ (μM) | Ligand<br>Efficiency<br>(kcal/mol) |
| --- | --- | --- | --- |
| 727590<br>(HTS hit) |  | 36* / 25.3** | 0.29* / 0.3** |
| Z56040660<br>(RNA primer<br>channel) |  | 5522* | 0.22 |
| Z26333448<br>(Thumb site II) |  | 2800** | 0.21 |
| RLA-6625<br>(Z285233820)<br>(Index–middle<br>finger pocket) |  | ~45%<br>(Inhibition at 10 mM) * | - |
| RLA-6626<br>(Active site,<br>R isomer of<br>Z190780124) |  | ~45%<br>(Inhibition at 10 mM) * | - |
| RLA-6627<br>(Active site,<br>S isomer of<br>Z190780124) |  | 6863* | 0.2 |
| RLA-6628<br>(derivative of<br>Z190780124) |  | 4905* | 0.19 |
| RLA-6629<br>(Active site) |  | ~30%<br>(Inhibition at 10 mM) * | - |

\* Determined with the gel-based assay; \*\*Determined with the PicoGreen assay

Given that the crystal structure of the complex with Active site hit Z190780124 suggested binding of the S enantiomer (Fig. 3A), we synthesized both stereoisomers and tested them in our inhibition assay (Fig. 8C, Table 2 and Fig. S12). Interestingly, the S isomer (RLA-6627) displayed more potent inhibition (IC_50_ value of 6.9 mM) than the R isomer (RLA-6626, 45% inhibition at 10 mM). The observed inhibition suggests thus a direct correlation between crystallographic binding and enzymatic interference for 4 of the 5 binding sites revealed by the fragment screening campaign, suggesting that these pockets represent functionally relevant regions within the 3D^pol^ architecture.

## DISCUSSION

Despite the near eradication of polio due to the development of the vaccine against poliovirus, there is a great need for broad-spectrum antiviral compounds to treat infections caused by other picornaviruses, such as enterovirus D68, enterovirus A71, poliovirus, coxsackievirus, and rhinovirus (54). RdRps play a pivotal role in picornavirus replication and exhibit significant sequence conservation; hence, RdRp-based drug design for treating picornavirus infection can be promising.

The structures obtained from this CFS campaign represent the highest-resolution EV-D68 3D^pol^ structures to date, with resolutions ranging from 1.34 to 1.83 Å. In all the structures, including the one in the ground state, we have observed that the loop at the entrance of the NTP channel is closed, similar to that observed in the GTP-bound structure of EV-D68 3Dpol (Fig. S7) (49).

The EV-D68 3D^pol^ has high sequence identity with other picornaviral RdRps: EV-A71 (64%), poliovirus (74%), coxsackievirus (74%), and human rhinovirus (56%), except for the foot-and-mouth disease virus (32%). The RNA template channel, Active site, and the RNA primer channel sites identified in this work are highly conserved (Fig. S2). Hence, fragment hits at these sites might be exploited to develop broad-spectrum drugs.

The Active site is the most conserved of all the sites, with fragment hit Z190780124 offering an “anchoring point” for development of a small molecule inhibitor, with the strong hydrogen bonding interaction with the side chain of the catalytic site D325 of the GDD motif (Fig. 3A). Owing to the high resolution of the structure, we surmised that the bound Z190780124 adopts the stereoisomer S configuration (Fig. 3A), which has been validated through the inhibition activity assays comparing both isomers (which we synthesized). Indeed, the S isomer displayed a substantially higher inhibition than the R isomer (Fig. 8, Table 2 and Suppl. Data). Picornaviral RdRps use the two-metal catalytic mechanism like other replicative polymerases. In the ground state structure and all the fragment-bound structures, we observed a Mg^2+^ ion at the active site (Fig. S8B); these are the first structures of EV-D68 3D^pol^ with metal ion at the active site.

Regarding the RNA template and primer channels, although we did not observe inhibitory activity for fragments binding to the RNA template channel, both sites partially overlap with the binding sites of several small-molecule inhibitors of human enterovirus and the related norovirus RdRps (Table S2). Indeed, these compounds could interfere with the positioning of the RNA duplex in the groove (Fig. 2F and 4H).

For the RNA template channel, the hits partially overlap with the abundant natural cofactor NADPH, bind the RNA template channel and inhibit EV-D68 3D^pol^ (15). Additionally, RNA template channel lead inhibitor GPC-N114, which also partially overlaps with the hits, exhibited antiviral activity against a panel of enteroviruses (EC_50_ values 0.1 - 2.0 µM, with EC_50 EV-D68_ of 1.44 µM) (39). The fact that pre-incubation of the inhibitors before the RNA template-primer duplex was necessary for inhibition (39,44) supports the notion that their interaction to 3D^pol^ affects binding of the template-primer to the RdRp. For the RNA template channel, previous literature also indicates that this site holds promise for inhibition of flaviviruses such as dengue virus (55–57).

For the RNA primer channel, the hits partially overlap with norovirus RdRp inhibitors (of the related *Caliciviridae* family): suramin-related compounds (43) and two naphthalene-sulfonate derivatives: NAF2 and PPNDS (Table S2). PPNDS displays an IC_50_ value of 0.45 µM against recombinant human norovirus RdRp (44). The RNA primer channel, apart from housing the primer strand, contains an important residue, L21, which is a part of the long loop of the finger domain that contributes significantly to the stability of the enzyme by bridging the finger and thumb domains together (49,58). Moreover, it is close to the NTP entry channel (Fig. S1B), essential for the access of NTPs for incorporation into the primer strand. From the strategic location of this site, we argue that inhibitors developed to target this site would have the potential to interfere with the addition of NTPs, apart from inhibiting RNA synthesis.

Regarding the two hotspot sites outside of the RNA binding groove, the inhibitory activity we show here (Fig. 8 and Table 2) with Z26333448 (Thumb site II, IC_50_ = 2.8 mM) and RLA-6625 (or Z285233820, Index-middle finger pocket, 45% inhibition at 10 mM) supports the notion that they affect the function of 3D^pol^. The Thumb site II seems to be coincidentally in an equivalent position to its flaviviral and hepaciviral counterparts, as previously mentioned (Fig. 5H-I). Nevertheless, its location at the base of the thumb subdomain may be relevant for the overall stability and dynamics of the enzyme, and small molecule binders might perturb these aspects.

In the case of the Index-middle finger pocket, it is in the vicinity of the ‘kink’ region identified in EV-A71 3D^pol^ elongation complex structure (Fig. S9A). The kink is a conserved region in picornaviral RdRps and has been speculated to optimize RdRp function through its ability to interact with RNA bases (59). There are no binders reported for this site in RdRps before and this opens possibilities of exploring the structural and functional significance of this new binding pocket. Modeling two additional 5’ template strand bases on the EV-A71 3D^pol^ elongation complex structure allows the last nucleobase to fit readily into the Index-middle finger pocket (Fig. S9B). Thus, we can hypothesize that this pocket may have a similar role to the kink, contributing to the processivity of the enzyme, as argued for the kink pocket (59). Also, some of the fragments in this pocket interact with I3 and N6, that are located on the long loop of the finger domain that bridges the finger and thumb domains together. Truncation of this loop affects the thermal stability of 3D^pol^ (49). Hence, we speculate that inhibitors targeting this pocket could affect the stability of the enzyme.

The abundance of hits localized within these five sites—especially the RNA primer channel, Thumb site II, and Index-middle finger pocket—suggests they are highly amenable to scaffold growth. This structural landscape provides thus an opportunity to leverage algorithmic design tools to combine adjacent fragments—a strategy successfully implemented in previous CFS campaigns (60–62). Such an approach offers a rational pathway for the computational evolution of these initial small molecules into more potent, multi-site inhibitors.

Additionally, the discovery of fragment hits in the Active site, including fragment hit Z190780124 S isomer that interacts with catalytic residue D325, offers fragment-linking prospects with either the RNA template channel or the RNA primer channel (Fig. 9A). Finally, the HTS has unveiled a hit-to-lead compound, 727590, that provides a good option for subsequent development, with favorable physicochemical properties (Fig. 9B). To note also that we have calculated the ligand efficiency for all hits allowing IC_50_ determination, including 727590. Ligand efficiency is a measure of binding affinity per non-hydrogen atom in the compound and is a useful metric in drug development (63,64). Interestingly, 727590 shows a ligand efficiency of 0.3 Kcal/mol/non-hydrogen atom, which is considered to bear potential to lead to potent drug-like compounds (63,64).

**Figure 9:**
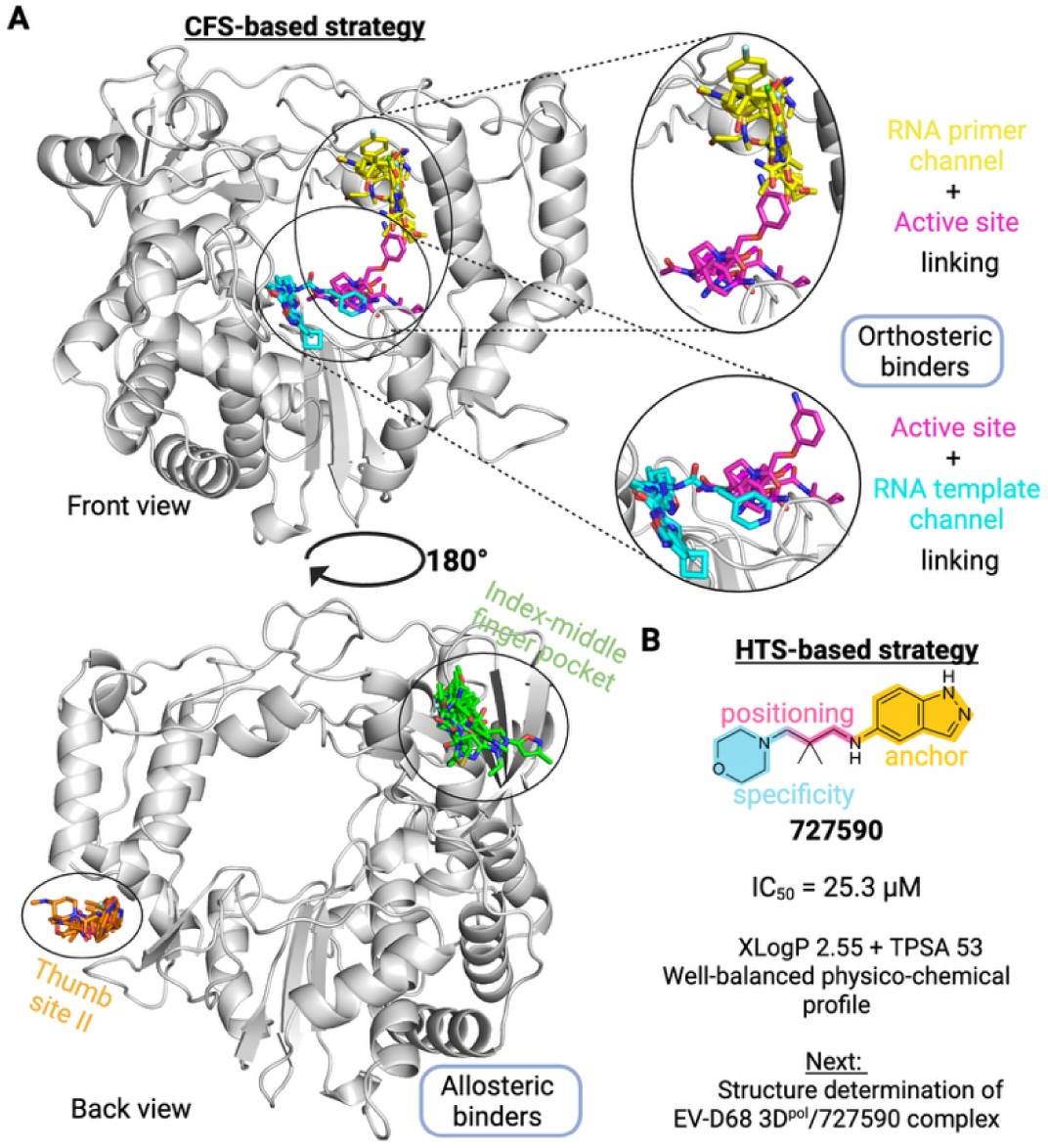
Prospects for design of EV-D68 3D^pol^ RdRp non-nucleoside inhibitors. A) CFS-based strategy. B) HTS-based strategy. Physico-chemical properties of 727590 calculated with DataWarrior v06.05.02. Created with biorender.com and the PyMOL Molecular Graphics System, version 2.5.0. Schrödinger, LLC.

Extensive SAR for this hit is beyond the scope of the present work, but the 5-aminoindazole core is established as the anchoring moiety and the morpholine group provides a specific size, shape and polarity determinant, both connected by a linker region that positions the previous moieties for engagement with EV-D68 3D^pol^. We are currently engaged in structure determination efforts for EV-D68 3D^pol^ in complex with the hit-to-lead compound 727590, which will provide a detailed SAR rationale and support subsequent development.

Together, these findings establish a framework for discovering non-nucleoside inhibitors of enteroviral RdRps and highlight the value of combining fragment-based and high-throughput approaches in antiviral drug discovery.

## Supporting information

Crystallography table

Supplementary Materials section

## ACKNOWLEDGEMENTS

We would like to acknowledge the Diamond Light Source for access to the fragment screening facility XChem (in particular, Blake Balcomb and Daren Fearon), for usage of DSi-Poised library, and for beamtime on beamline I04-1 under proposal LB38067.

## AUTHOR CONTRIBUTIONS

I.B. and Q.W. contributed as first authors equally to this work. F.X.R., E.A., M.A., and R.J.N. conceived the study and provided supervision. I.B. carried out protein expression, purification, crystallization, and diffraction data collection with assistance from M.S. I.B. and F.X.R. analyzed and processed the crystallographic data, with resources provided by E.A. Q.W. established the high-throughput biochemical screening assay, with assistance from L.N., E.P.T., J.L.R., with resources provided by M.G., R.J.N. and M.A. J.M. and J.C synthesized fragments with resources and supervision provided by A.R.R. E.P.T. established the gel-based primer extension assay, with resources provided by M.G. I.B., E.A., and F.X.R. wrote the initial draft, with assistance from Q.W. and J.M., and reviewed and edited by all the authors.

## SUPPLEMENTARY DATA

Supplementary Data are available at NAR online.

## DATA AVAILABILITY

The coordinates and structure factors have been deposited in the Protein Data Bank, under group deposition ID G_1002366 for the ligand-bound datasets, and under PDB 14AA for the ground dataset. All the other data needed to evaluate the conclusions in the paper are present in the paper and/or the Supplementary Data.

