## Supplementary Materials section for "Discovery of non-nucleoside inhibitors of the enterovirus D68 3D polymerase through crystallographic fragment and high-throughput biochemical screening"

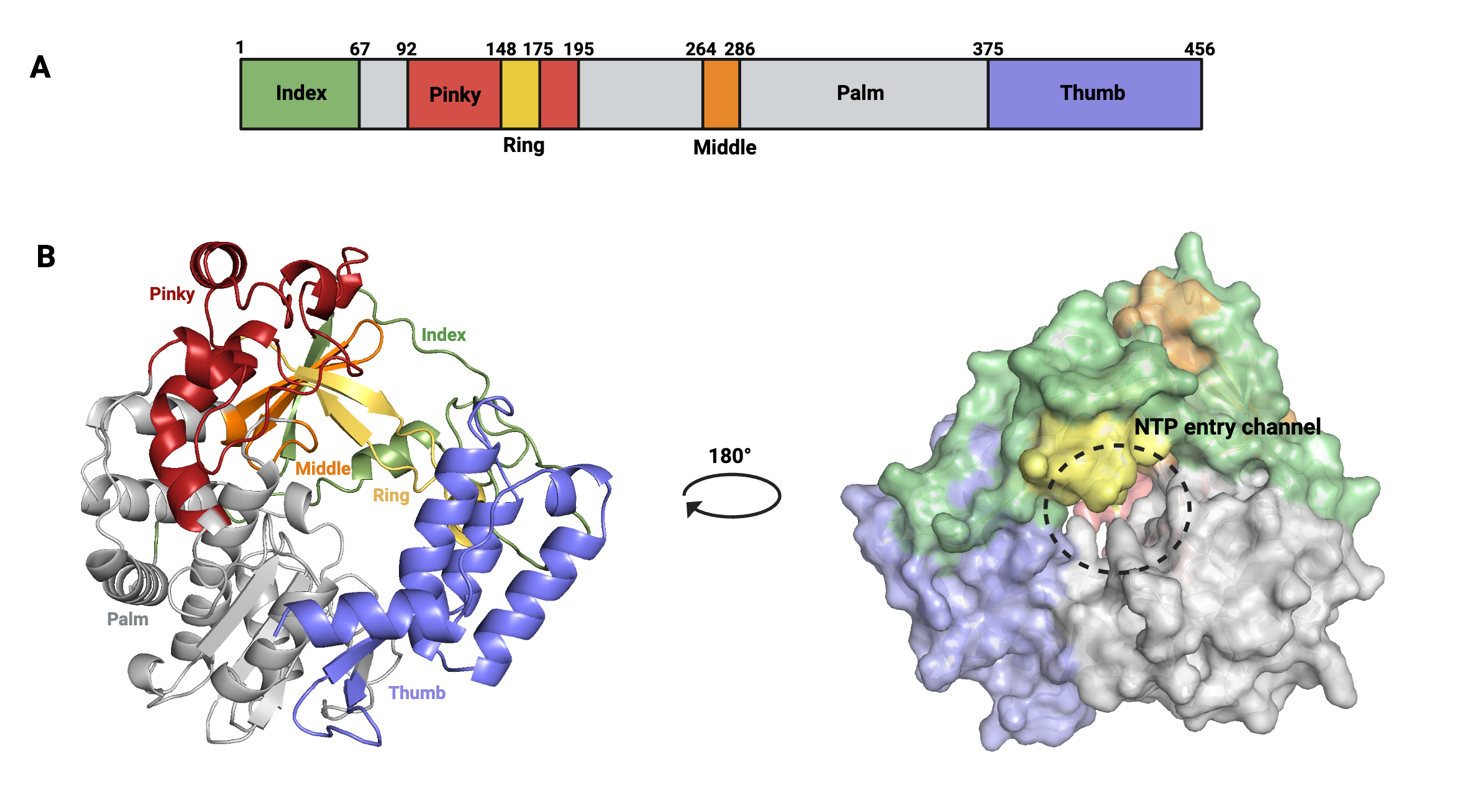

**Figure S1:** A) Domain organization of EV-D68 3D^pol^, each domain is color coded. B) Overall structure of EV-D68 3D^pol^ with the different domains labeled with the corresponding color (left), surface representation of the structure at 180° rotation, with the opening to the NTP entry channel marked (right).

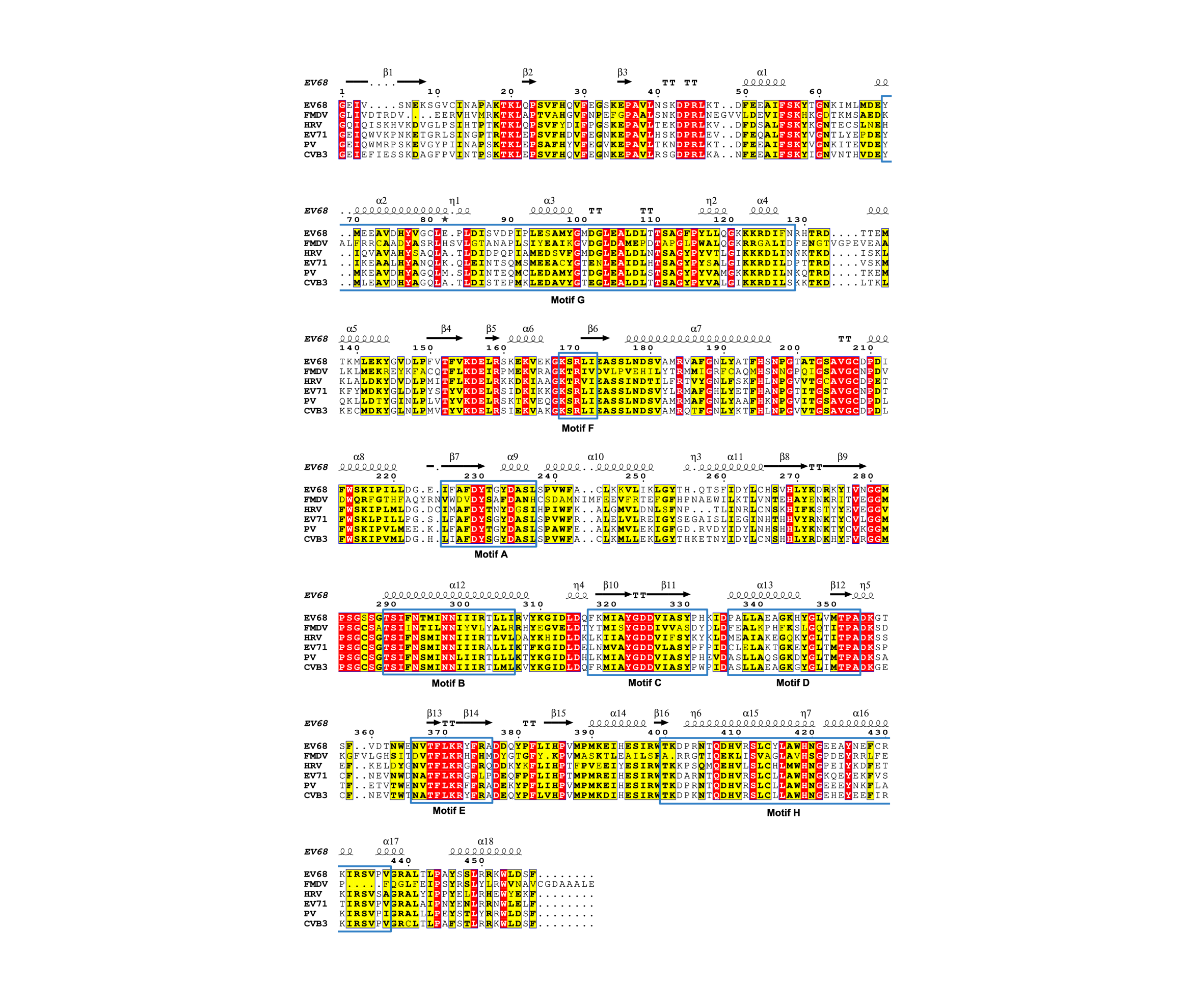

**Figure S2:** Multiple sequence alignment of RdRPs from the picornavirus family. The secondary structural elements of EV-D68 3Dpol are marked on top the alignment. The conserved structural motifs of picornaviral RdRps (A–H), are highlighted in box. Strictly identical residues are highlighted with a solid red background and white text, while highly conserved and similar residues are highlighted with a yellow background. Secondary structure elements extracted from [PDB ID / DSSP] are displayed above the alignment block: helices are represented by squiggles (α: α-helices; 10: 3₁₀-helices; η: hairpin loops, π: π-helices), β-strands are rendered as arrows, and strict turns are labeled as TT (β-turns) or TTT (α-turns). Numbers above the secondary structure elements indicate the position relative to the reference sequence.

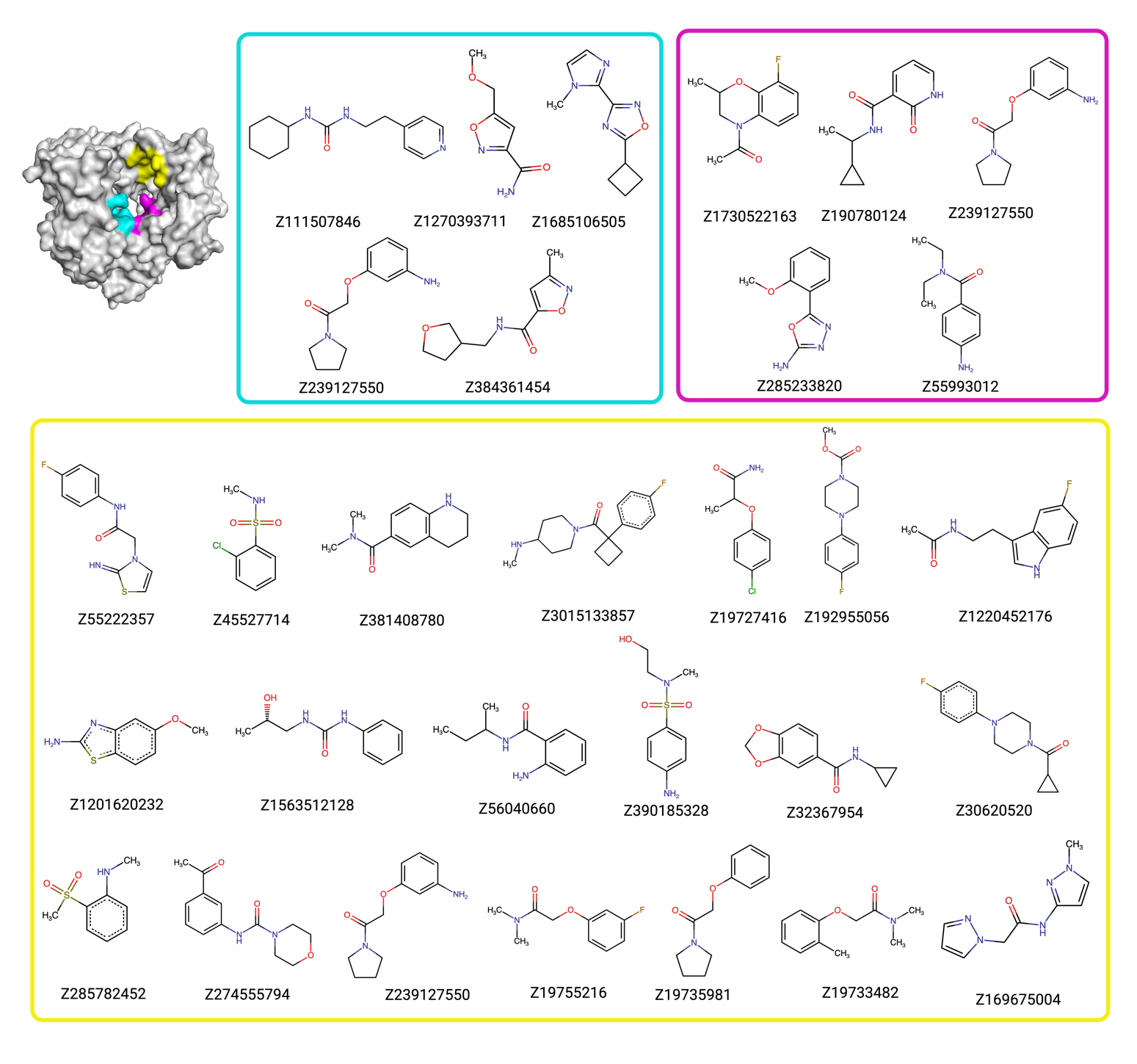

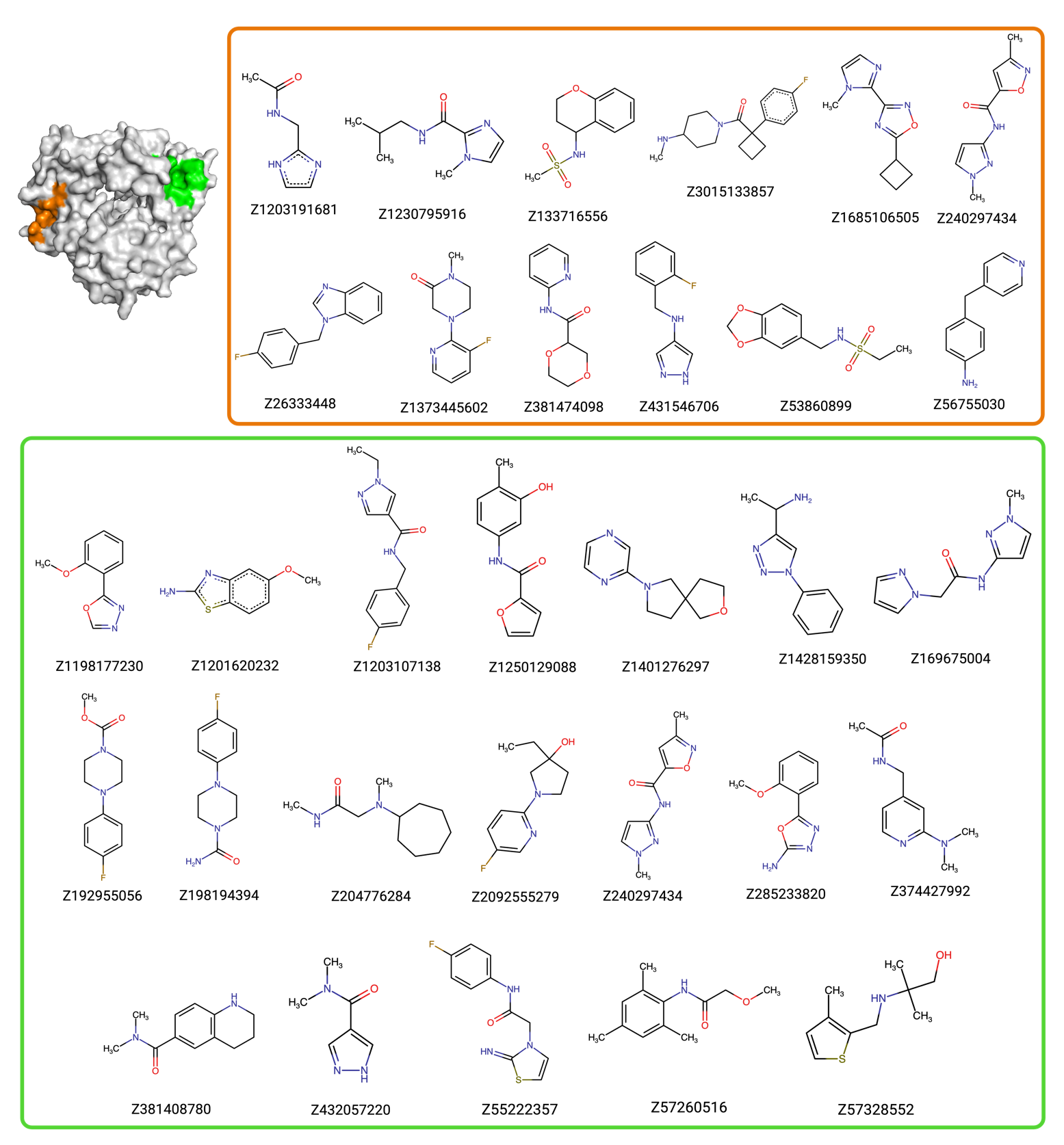

**Figure S3:** Full list of the fragment hits for the five binding pockets. Blue box: RNA template channel; Pink box: Active site; Yellow box: RNA primer channel; Orange box: Thumb site II; Green box: Index-middle finger pocket.

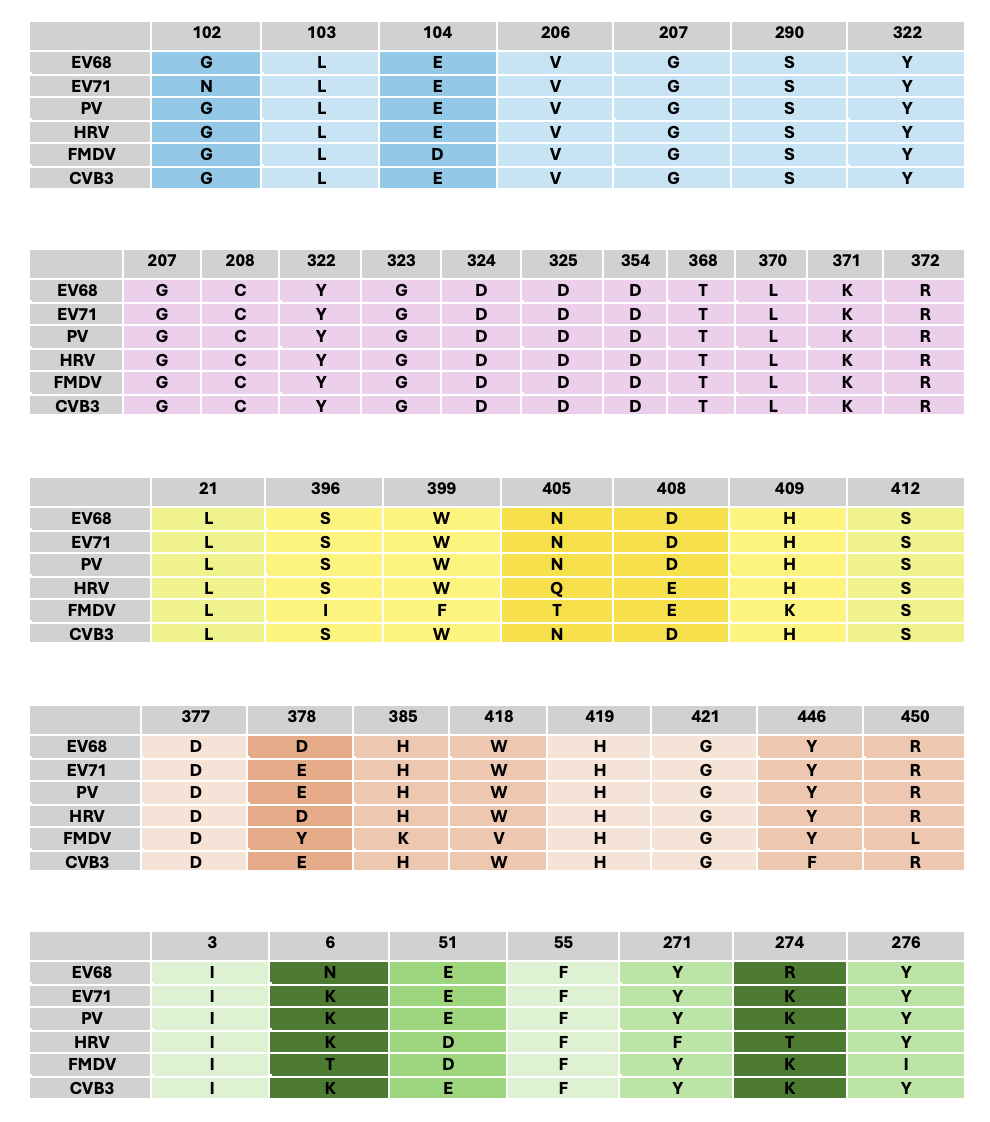
**Figure S4:** The sequence conservation of residues at the RNA template channel (blue), Active site (pink), RNA primer channel site (yellow), Thumb site II (orange), and Index-middle finger pocket (green), across different members of the picornavirus family. Positions are colored with a darker shade depending on the number of residues at the corresponding position that are different.

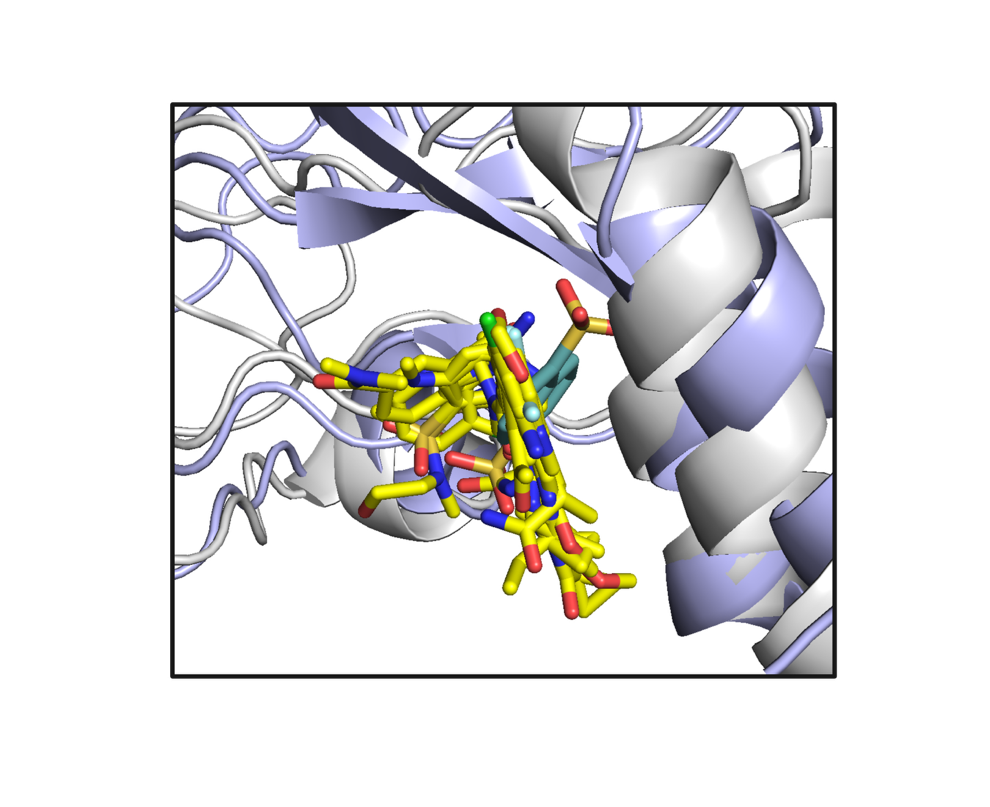

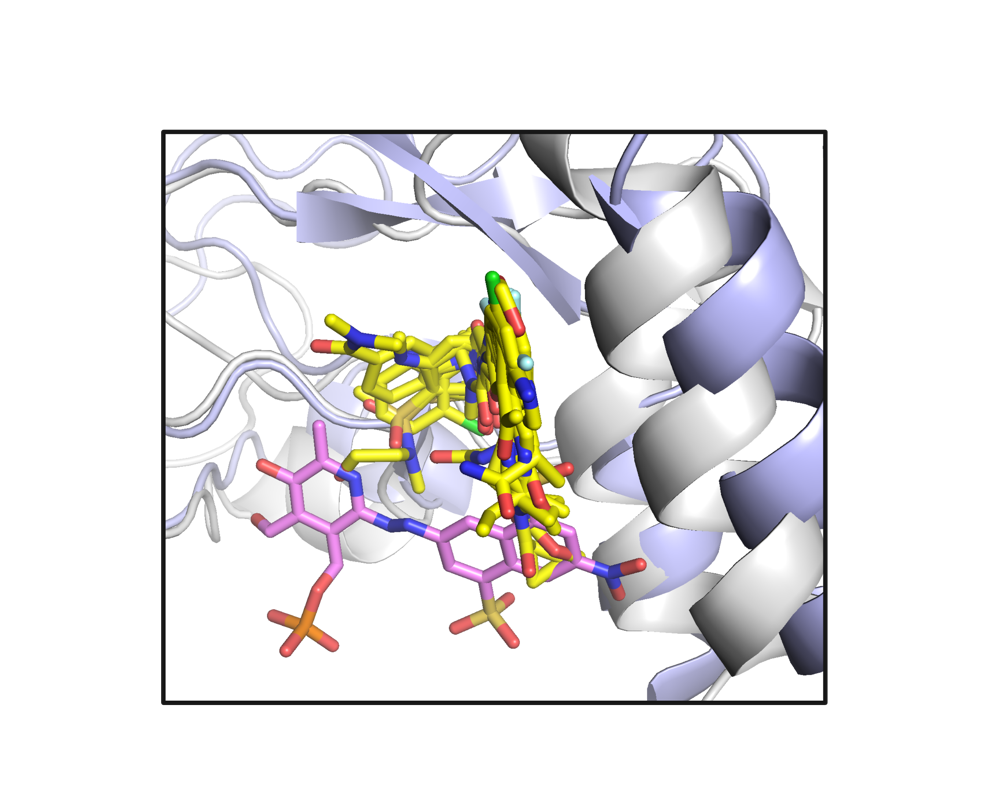

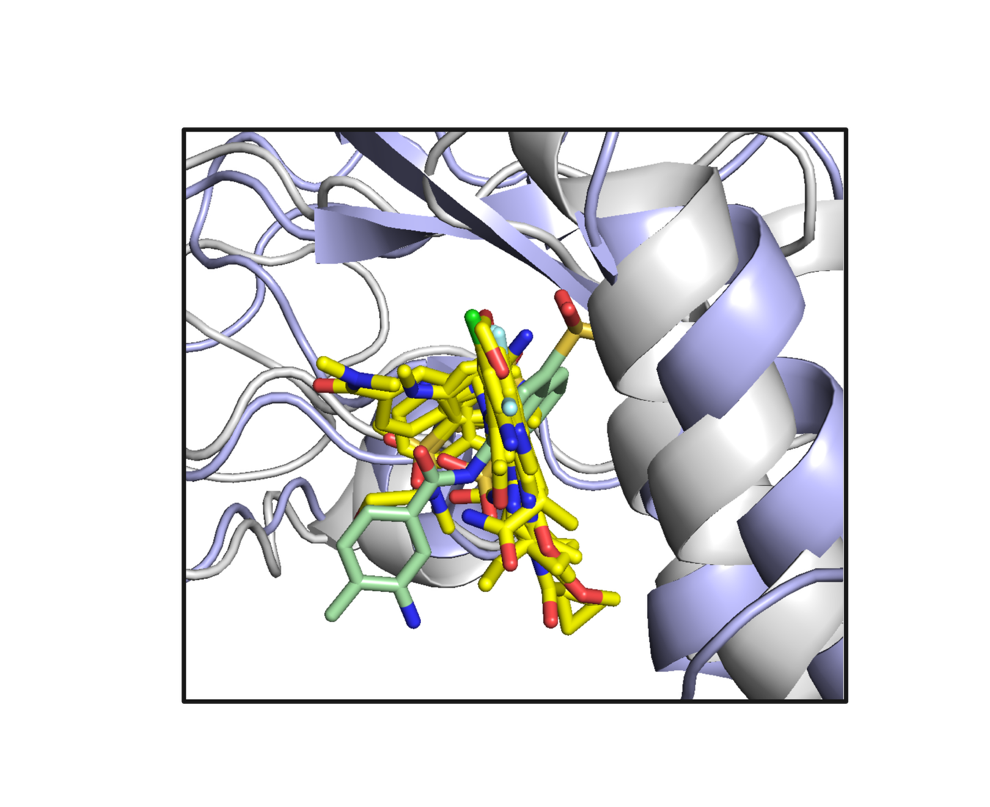

**B**

**A**

**C**

**Figure S5:** Superposition of fragment bound structures of EV-D68 3D^pol^ (grey) to human norovirus (hNV) RdRP (violet) bound to the small molecule inhibitors: A) Pyridoxal-5′-phosphate-6-(2′-naphthylazo-6′-nitro-4′,8′-disulfonate) tetrasodium salt (PPNDS) (pink) showing partial overlap between the binding sites (4LQ3); B) A suramin derivative compound (light green) showing partial overlap between the binding sites (4NRT); C) Naphthalene di-sulfonate (NAF2) (blue) showing complete overlap between the binding sites (4LQ9). The fragments at the RNA primer channel are shown in yellow.

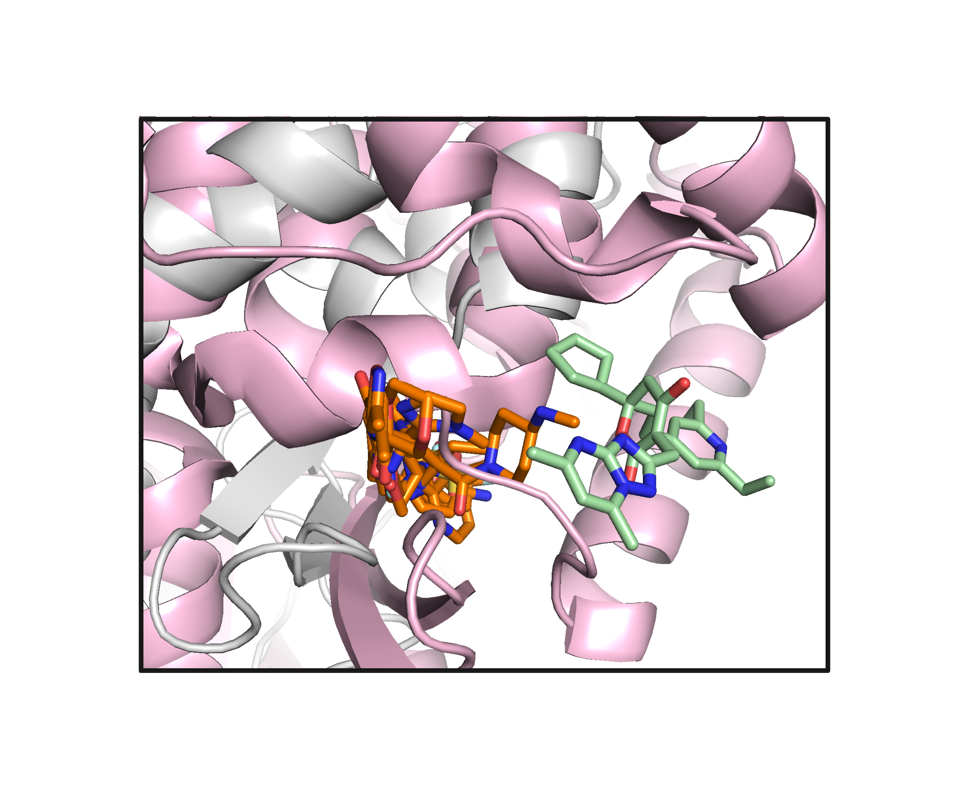

**Figure S6:** Superposition of fragment-bound structures of EV-D68 3D^pol^ (grey) to hepatitis C (HCV) RdRP (pink) bound to filibuvir (green) showing that the topology of the binding sites is different (3FRZ). The fragments at the Thumb site II are shown in orange.

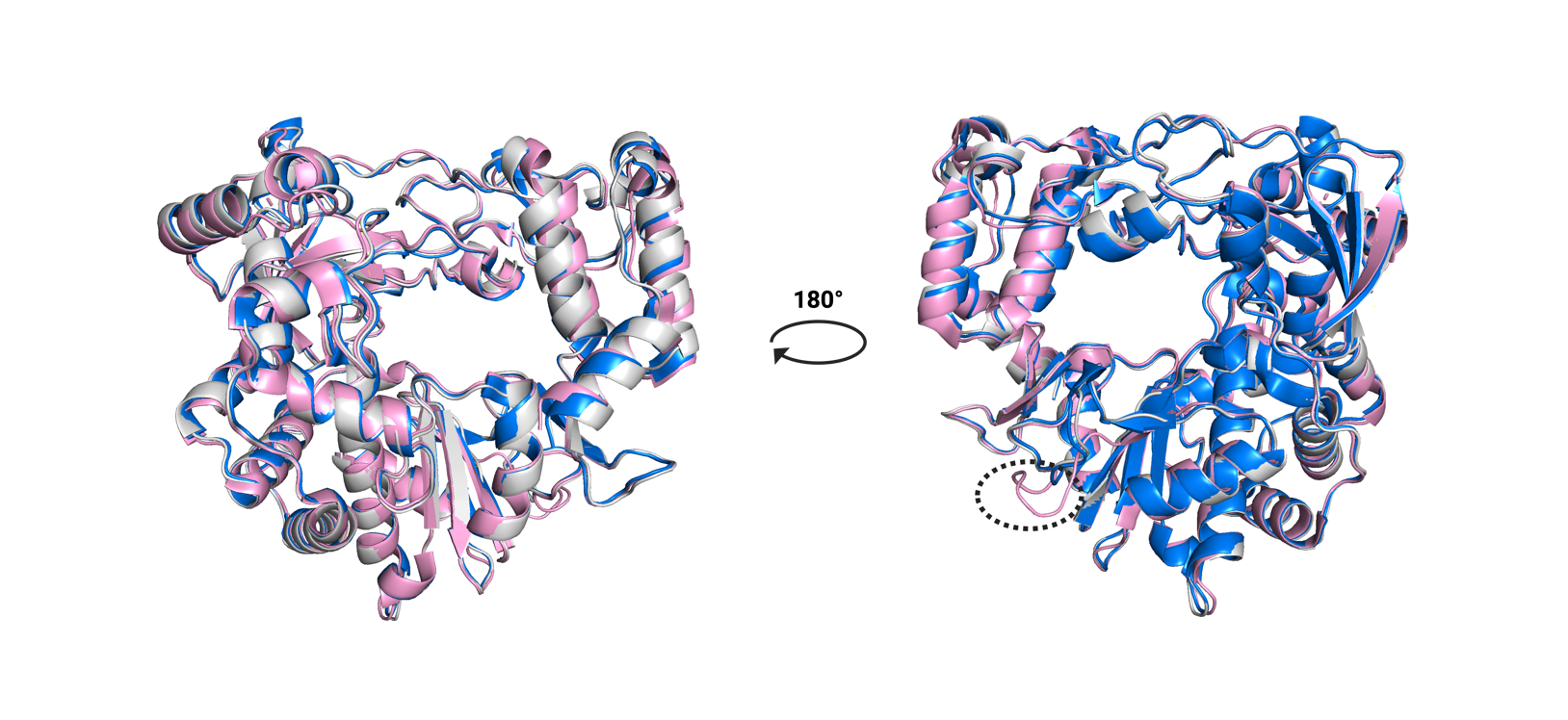
**Figure S7:** Superposition of the structure of EV-D68 3D^pol^ in ground state (grey) with the existing structures of EV-D68 3D^pol^: GTP bound structure (5XE0) shown in blue, and apo structure (6L4R) shown in pink. The loop spanning residues 351-362 is open in the apo structure but closed in our structure and the GTP-bound structures.

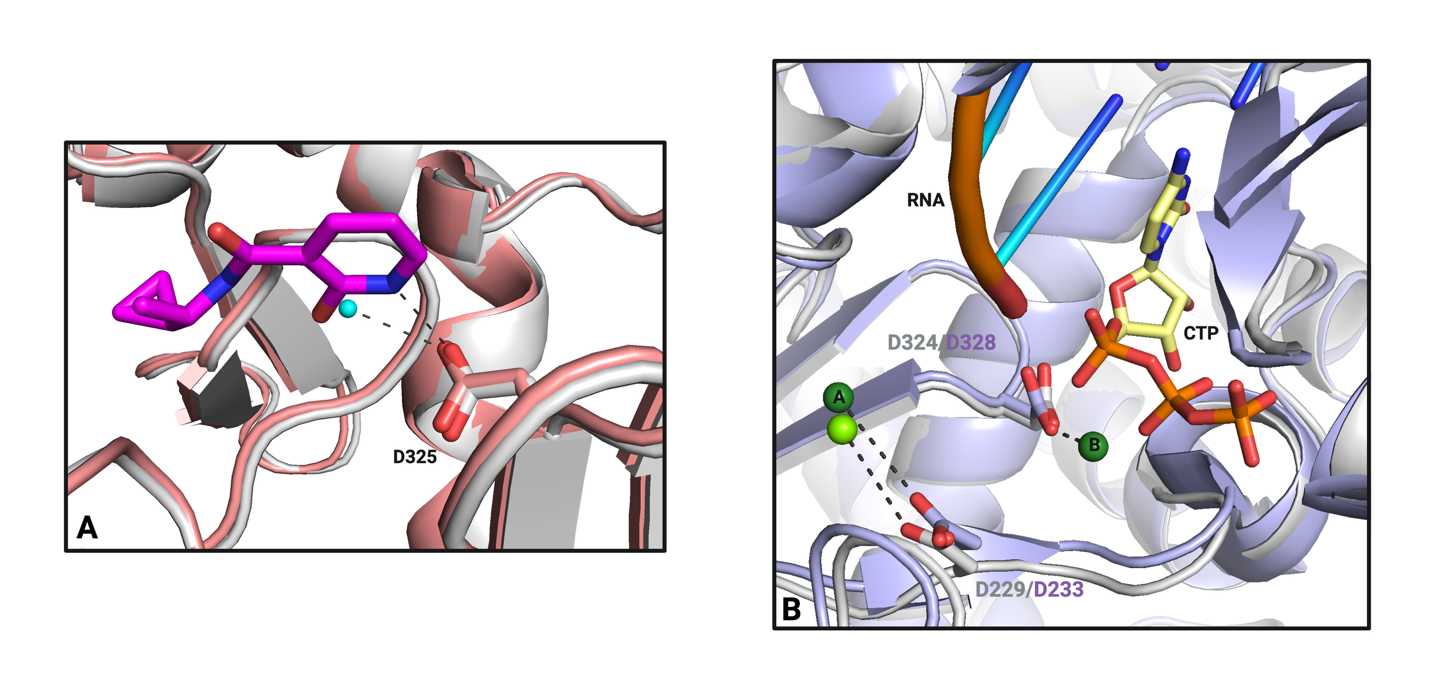

**Figure S8:** A) Superposition of fragment Z190780124 (magenta) bound EV-D68 3D^pol^ (grey) with ground state EV-D68 3D^pol^ (pink), displaying the water molecule (blue) interacting with the catalytic aspartate in the ground state which is replaced by Z190780124 in the fragment bound structure. B) Superposition of Apo EV-D68 3D^pol^ (grey) displaying the Mg^2+^ ion (light green) at the active site and, EV71 RdRp (violet) elongation complex with the active site in an open conformation (5F8I) displaying Mg^2+^ ions A & B (dark green).

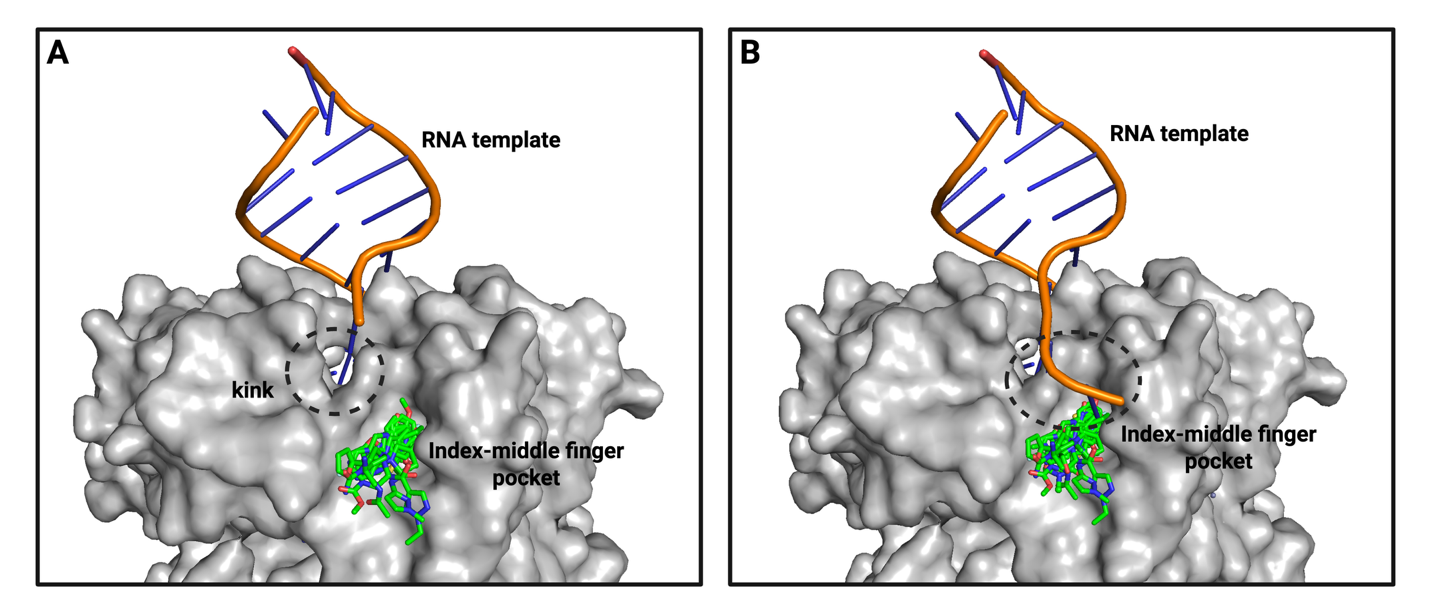

**Figure S9:** Superposition of the structure of EV-D68 3D^pol^ in ground state (grey) onto the EV-A71 3D^pol^ EC (6KWR), with the RNA template displayed (orange), the kink region on EV-D68 3D^pol^ marked (dashed circle) and the fragments bound to the Index-middle finger pocket (green).

**Table S2:** Two-dimensional structures and PubChem IDs of known inhibitors.

| **Small Molecule** | **PubChem ID** | **Structure** |
| --- | --- | --- |
| **NADPH**  **(RNA template channel)** | 5884 | 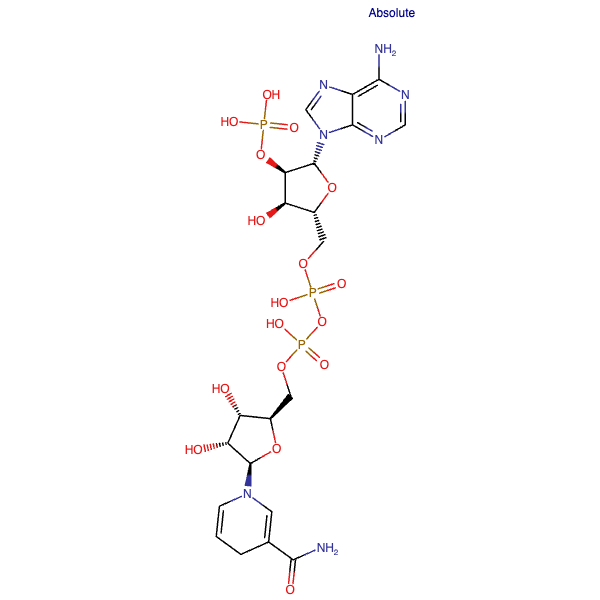 |
| **GPC-N114**  **(RNA template channel)** | 91666467 | 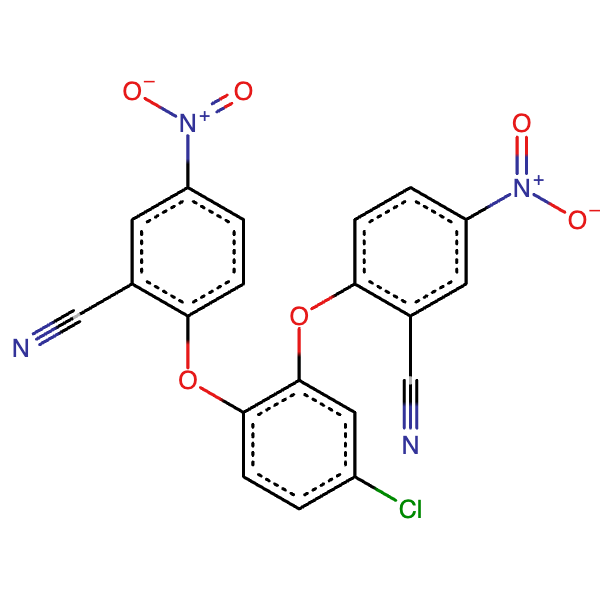 |
| **GPC-N143**  **(RNA template channel)** | 91666468 | 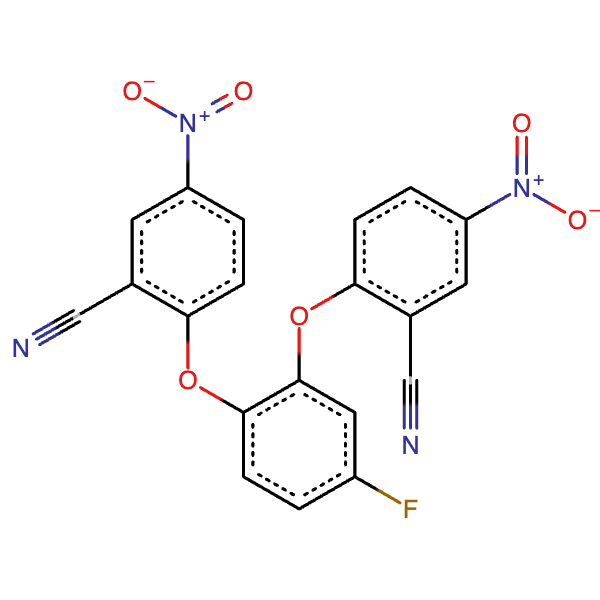 |
| **NAF2**  **(RNA primer channel)** | 6666 | 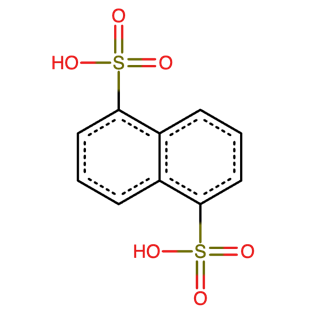 |
| **PPNDS**  **(RNA primer channel)** | 3932023 | 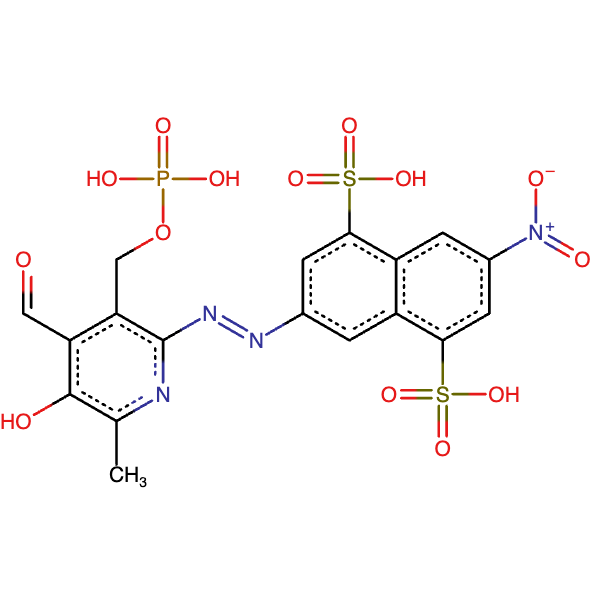 |
| **Filibuvir**  **(Thumb site II)** | 54708673 | 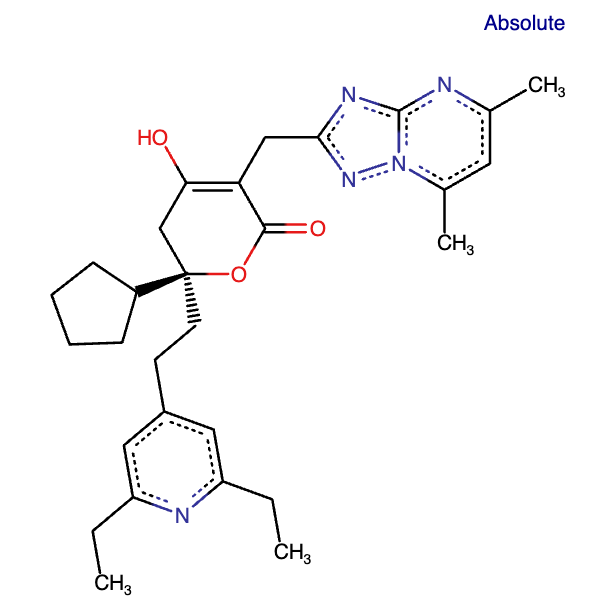 |
| **Lomibuvir**  **(Thumb site II)** | 24798764 | 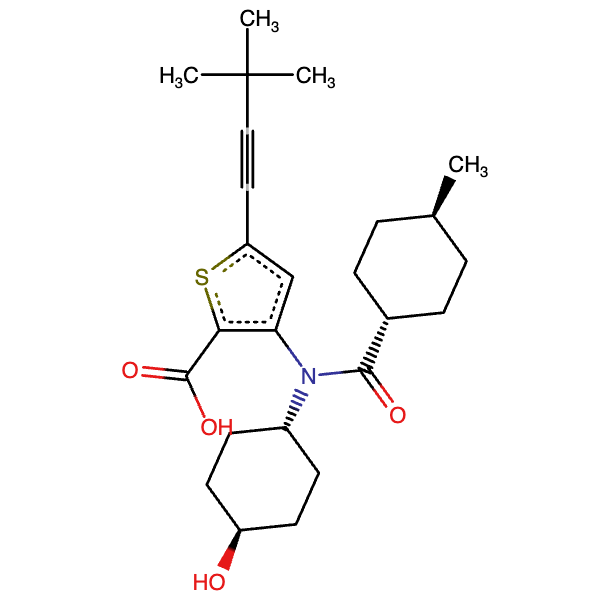 |
| **Radalbuvir**  **(Thumb site II)** | 53259022 | 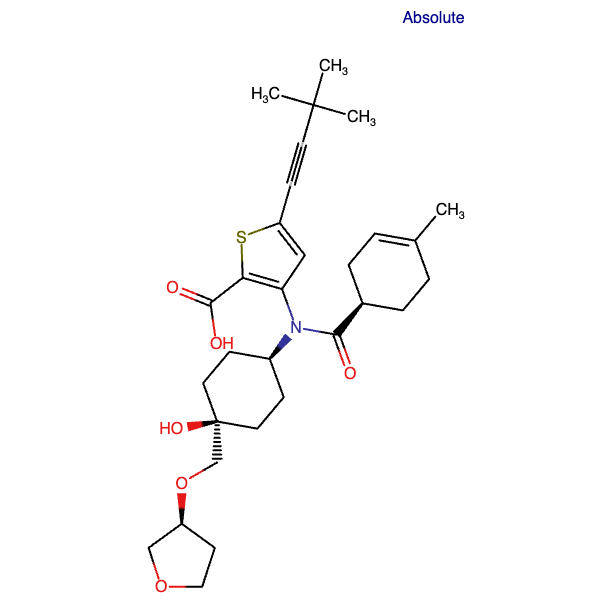 |

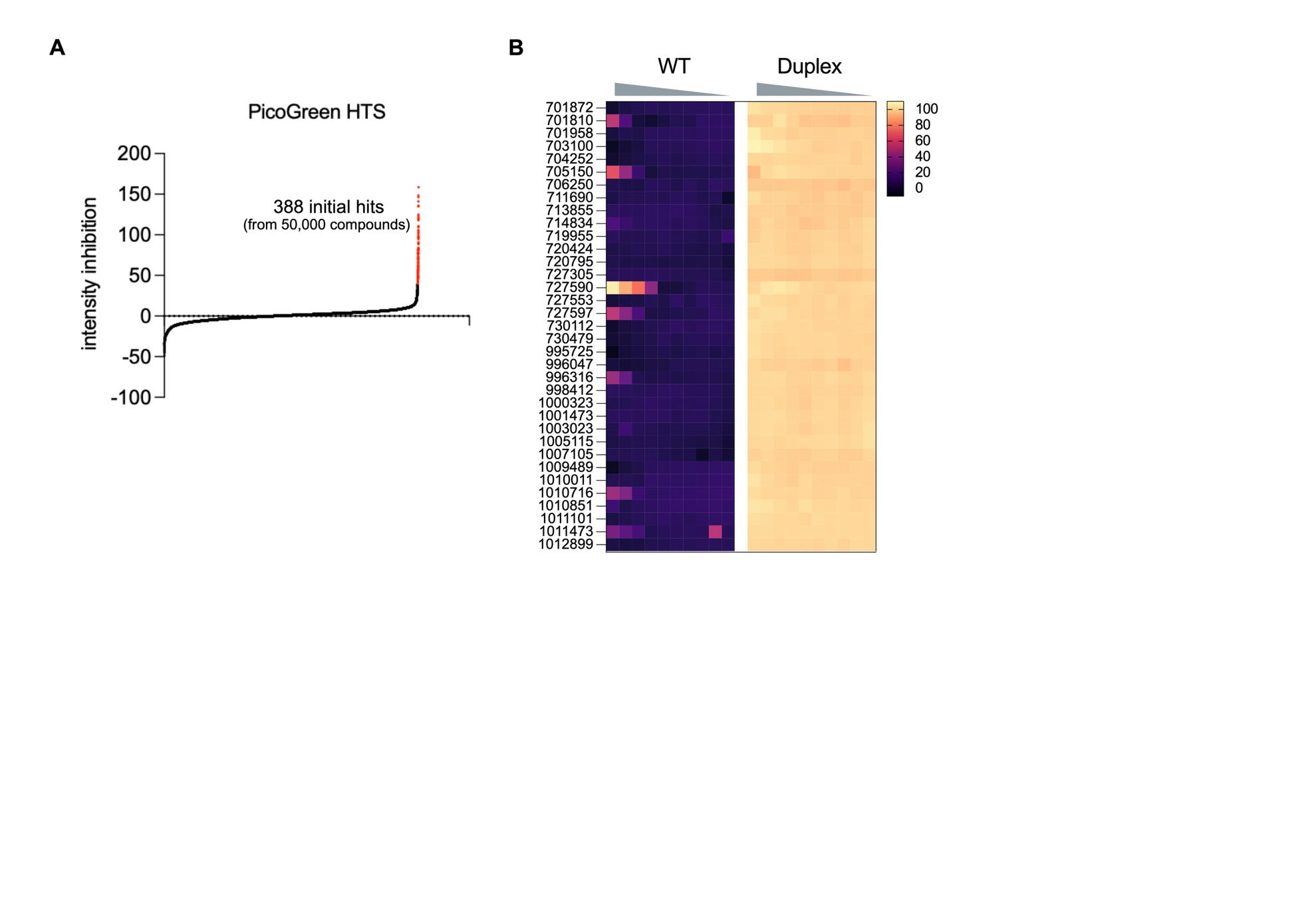

**Figure S10: Initial hit determination and counterscreening with selected 35 hit compounds:** For detail of the PicoGreen assay, the reader is referred to the MATERIALS AND METHODS section in the main manuscript. A) 388 initial hits were identified (a "hit" was defined as a data point that lies more than three standard deviations away from the mean); B) The result of dose-responsive PicoGreen assay for the selected 35 compounds from the initial hits. In short, “WT” corresponds to the assay in presence of all the components, while the “Duplex” counterscreeen is performed with all the components except EV-D68 3D^pol^. Under “WT” and “Duplex”, a triangular wedge denotes the direction of increasing concentration of compound (from right to left). The heatmap coloring refers to inhibition percentage.

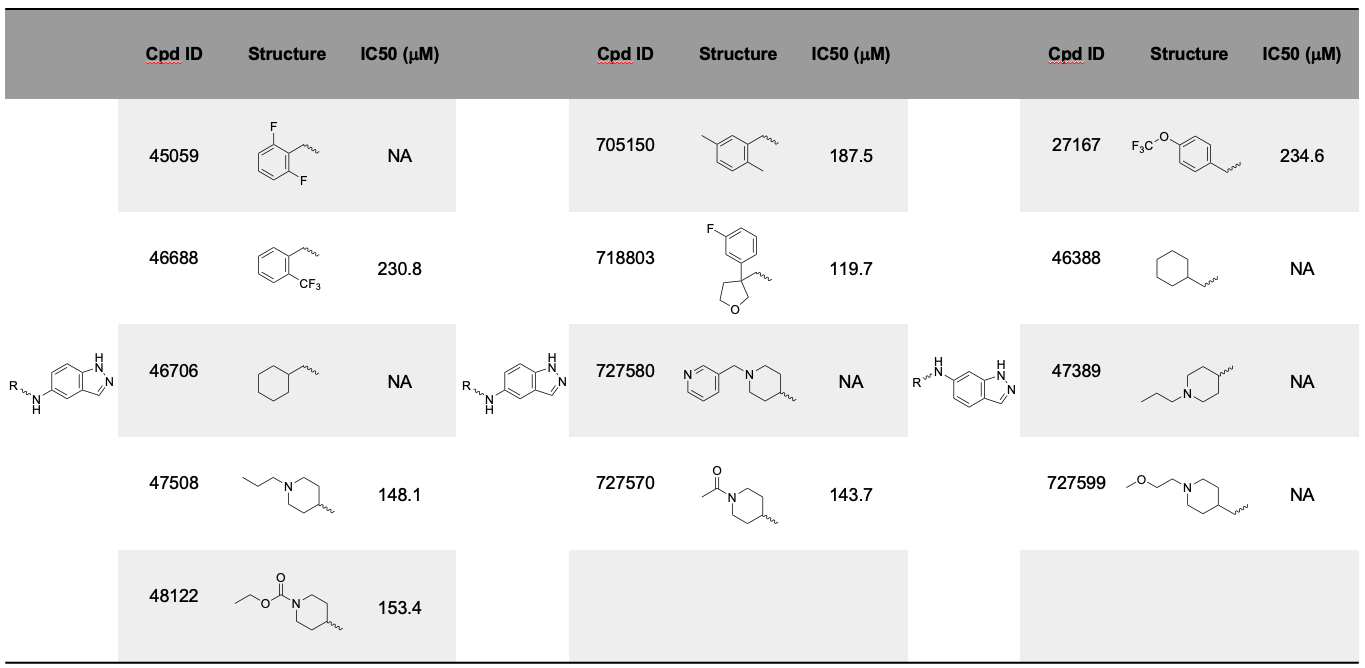
**Table S3.** Structures and IC_50_ values of thirteen additional compounds with either a 5-aminoindazole or a 6-aminoindazole core from the ChemBridge Premium and ChemBridge Gallo libraries.

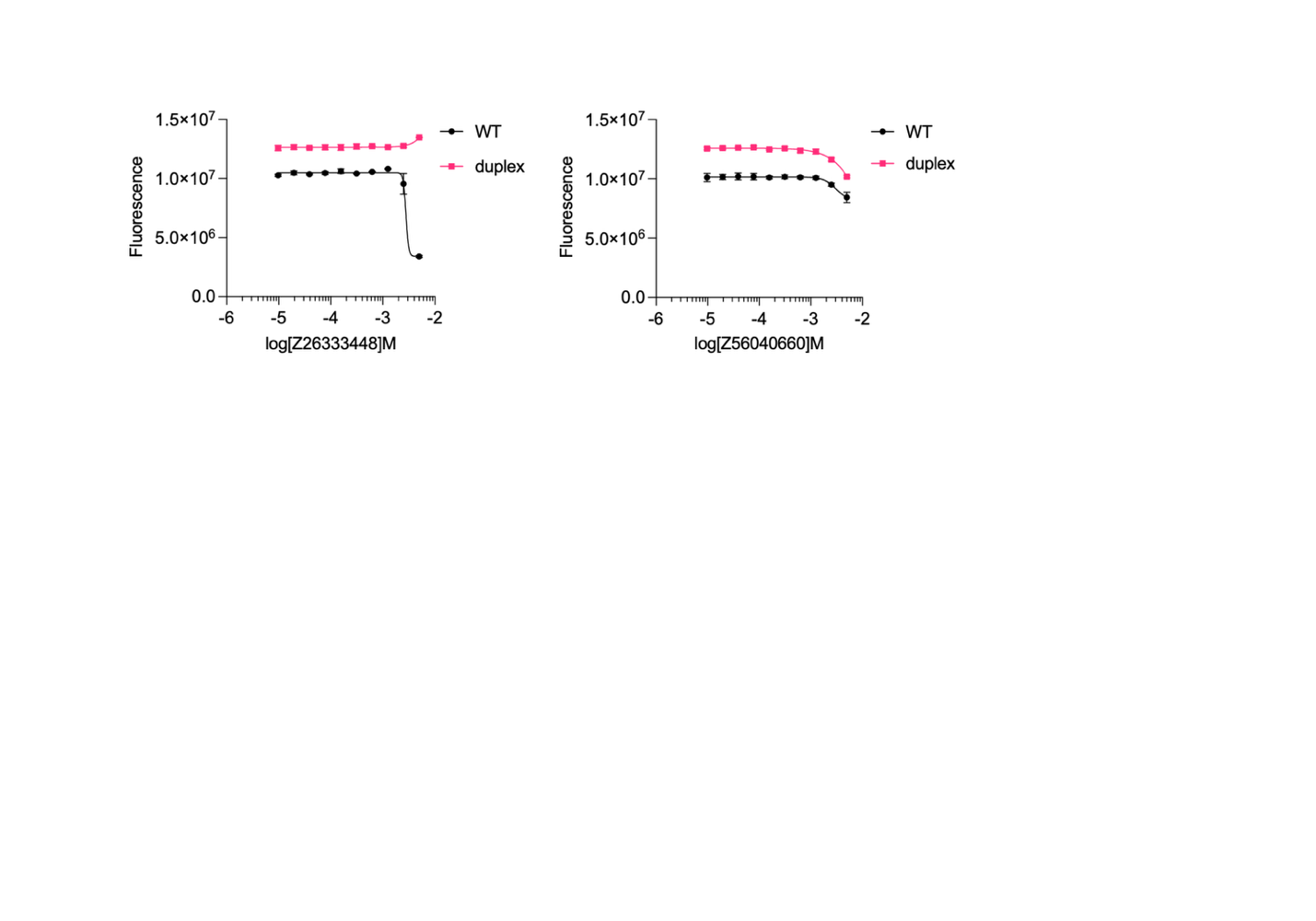

**Figure S11:** Counterscreen for compounds Z26333448 and Z56040660. In short, “WT” corresponds to the assay in presence of all the components, while the “Duplex” counterscreeen is performed with all the components except EV-D68 3D^pol^.

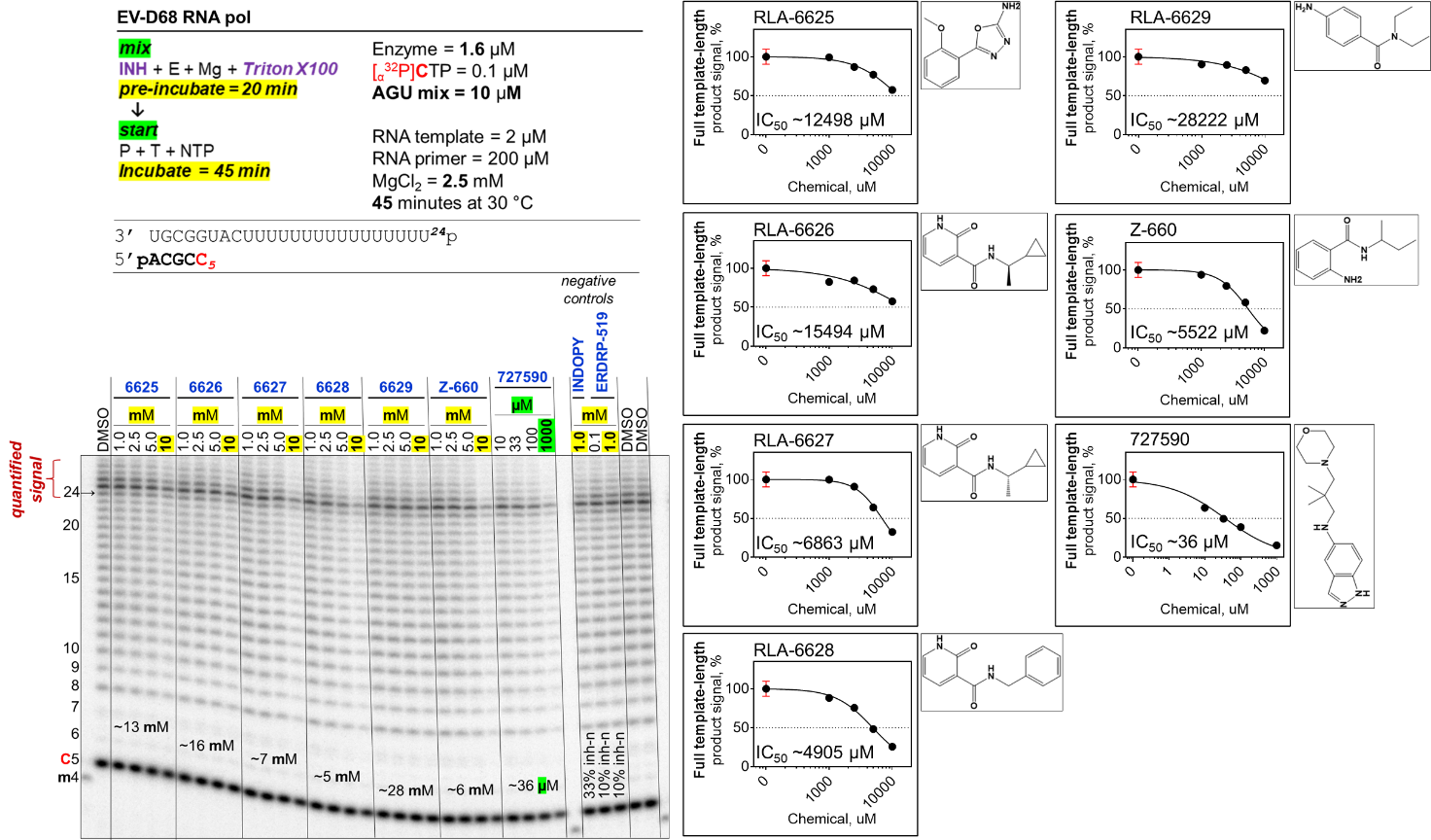

**Figure S12: Gel-based RNA primer extension assay results.** Top left: RNA synthesis (primer extension) assay conditions. Bottom left: representative annotated gel image readout of the primer extension assay illustrating the migration patterns of products of RNA synthesis on a denaturing 8M urea 20% polyacrylamide gel subjected to electrophoresis. Right: IC_50_ curves and values, with the chemical structure of each compound shown to the right of the corresponding curve.

**Chemistry**

Chemical reagents and anhydrous solvents were purchased from commercial suppliers and used without further purification. 4-amino-*N,N*-diethylbenzamide was purchased from Ambeed Inc. Thin layer chromatography (TLC) (silica gel, F254, 250 µm) were performed on precoated TLC glass plates and were visualized by fluorescence quenching under UV light. Chromatography was carried out using Isolera Four and CombiFlash NextGen 300 flash chromatography systems with SiliaSep silica gel from Silicycle. Chromatography used HPLC grade Hexanes and HPLC grade Ethyl Acetate. NMR spectra were recorded on a Bruker Advance III HD 400 MHz spectrometer. Chemical shifts (δ) are expressed in parts per million (ppm) and are referenced to CDCl_3_ (7.26 ppm) and DMSO-d_6_ (2.50 ppm). Coupling constants are reported as Hertz (Hz). Splitting patterns are indicated as follows: s, singlet; d, doublet; t, triplet; dd, doublet of doublets; qd, quartet of doublets; m, multiplet; br, broad singlet. Low-resolution ESI Mass Spectrometry was performed on a Waters Acquity UPLC QDa mass spectrometer equipped with Quaternary Solvent Manager, Photodiode Array Detector and Evaporative Light Scattering Detector. Separations were carried out with Acquity UPLCÒ BEH C18 1.7 μm, 2.1 x 50 mm column at ambient temperature. The mobile phase was MilliQ-H2O with 0.1% trifluoroacetic acid (eluent A) and HPLC grade MeCN with 0.1% trifluoroacetic acid (eluent B). Signals were monitored at 254 and 280 over 3.5 min, with a gradient of 5 to 95% eluent B for 2.75 min, then held at 100% B for 0.5 min.

**5-(2-methoxyphenyl)-1,3,4-oxadiazol-2-amine**

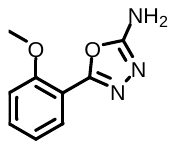

To a 10 mL flask was added semicarbazide HCl (164 mg, 1.47 mmol, 1.0 eq.) and sodium acetate (127 mg, 1.54 mmol, 1.05 eq.). Dissolve in DI water (2 mL) , then cool to 0 °C. Add 2-methoxybenzaldehyde (200 mg, 1.47 mmol, 1.0 eq.) dissolved in THF (1 mL), stir at 0 °C for 1.5 hr. After 1.5 hr, add more THF (2.5 mL, total 0.35 M to 2-methoxybenzaldehyde), followed by K_2_CO_3_ (508 mg, 3.67 mmol, 2.5 eq.) and Chloramine T trihydrate (538 mg, 1.91 mmol, 1.3 eq.) added as solids at 0 °C. Stir for 5 min at 0 °C, then remove from ice bath and stir at room temperature for 3 hr. After 3 hr., the crude solution is dumped into a separatory funnel, washed with a 1:1 mixture of sat. Na_2_S_2_O_3_ soln. and sat. NaHCO_3_ soln. and extracted with Toluene (3X). The organic layers were combined, then washed with 2M HCl soln. The aqueous layer was collected and added to a 50 mL flask, which was then adjusted to a pH >12 by slow addition of 10M NaOH soln. This was stirred at room temperature for 2 hr., after which the soln. was filtered, and the precipitate was washed with water. The filter cake was collected and dried to yield pure **5-(2-methoxyphenyl)-1,3,4-oxadiazol-2-amine** (197 mg, 1.03 mmol, 70 % yield) as a white powder.

Chemical Formula: C_9_H_9_N_3_O_2_; MW: 191.19.

LC/MS (ESI): m/z = 192.14 [M+H]^+^. Retention time on LC/MS: 1.74 min.

^1^H NMR (400 MHz, DMSO-*d*_6_) δ 7.70 (d, *J* = 8.1 Hz, 2H), 7.36 (d, *J* = 8.0 Hz, 2H), 7.26 (s, 2H), 2.37 (s, 3H).

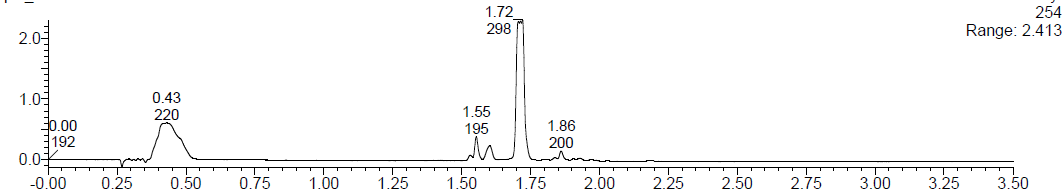

**N-[(1*R*)-1-cyclopropylethyl]-2-oxo-1H-pyridine-3-carboxamide**

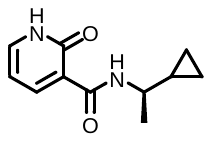

To a 20 mL vial was added 2-methoxynicotinic acid (400 mg, 2.61 mmol, 1.0 eq.) and HATU (1.04 g, 2.74 mmol, 1.05 eq). Dry MeCN (2 mL, 1.4 M to 2-methoxynicotinic acid) and NEt_3_ (1.09 mL, 7.84 mmol, 2.20 eq.) were then added, and the reaction was stirred at room temperature for 15 min. After, (*R*)-1-cyclopropylethan-1-amine HCl (318 mg, 2.61 mmol, 1.0 eq) was added to the vial and the reaction was allowed to stir overnight at room temperature. The next morning, the reaction was quenched into 1M HCl and extracted with CH_2_Cl_2_ (3X). The organics were combined, dried over Na_2_SO_4_, and concentrated *in vacuo*. The crude was then purified via silica gel chromatography (0 to 50% EtOAc in Hexanes, 15 CV) to yield (*R*)-N-(1-cyclopropylethyl)-2-methoxynicotinamide (404.1 mg, 1.86 mmol, 70% yield) as a beige solid.

(*R*)-N-(1-cyclopropylethyl)-2-methoxynicotinamide (269.0 mg, 1.22 mmol, 1.0 eq) was added to a 20 mL vial along with 4M HCl in Dioxane (10.2 mL, 0.12 M to (*R*)-N-(1-cyclopropylethyl)-2-methoxynicotinamide). The reaction was brought up to 90 °C and stirred overnight. After the overnight, the solvent was removed, the crude was suspended in MeOH then filtered. The precipitate was then purified via silica gel chromatography (0 to 10% MeOH in CH_2_Cl_2_, 15 CV, dry loaded with celite) to yield **N-[(1*R*)-1-cyclopropylethyl]-2-oxo-1H-pyridine-3-carboxamide** (127 mg, 0.62 mmol, 50% yield) as a crystalline, white solid.

^1^H NMR (400 MHz, DMSO-*d*_6_) δ 12.46 (br, 1H), 9.79 (d, *J* = 8.0 Hz, 1H), 8.31 (dd, *J* = 7.1, 2.3 Hz, 1H), 7.69 (dd, *J* = 6.2, 2.3 Hz, 1H), 6.46 (dd, *J* = 7.1, 6.2 Hz, 1H), 3.49 (qd, *J* = 8.0, 6.6 Hz, 1H), 1.19 (d, *J* = 6.6 Hz, 3H), 1.00 – 0.87 (m, 1H), 0.48 – 0.35 (m, 2H), 0.23 (dd, *J* = 16.8, 3.6 Hz, 2H).

Chemical Formula: C_11_H_14_N_2_O_2_; MW: 206.24.

LC/MS (ESI): m/z = 207.21 [M+H]^+^. Retention time on LC/MS: 1.66 min.

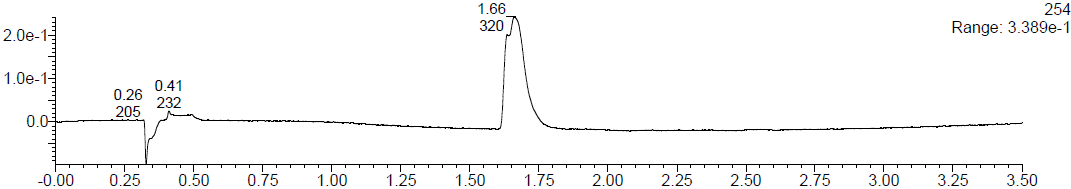

**N-[(1*S*)-1-cyclopropylethyl]-2-oxo-1H-pyridine-3-carboxamide**

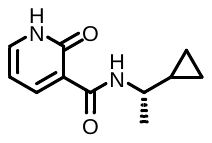

To a 20 mL vial was added 2-methoxynicotinic acid (400 mg, 2.61 mmol, 1.0 eq.) and HATU (1.04 g, 2.74 mmol, 1.05 eq). Dry MeCN (2 mL, 1.4 M to 2-methoxynicotinic acid) and NEt_3_ (1.09 mL, 7.84 mmol, 2.20 eq.) were then added, and the reaction was stirred at room temperature for 15 min. After, (*S*)-1-cyclopropylethan-1-amine HCl (318 mg, 2.61 mmol, 1.0 eq) was added to the vial and the reaction was allowed to stir overnight at room temperature. The next morning, the reaction was quenched into 1M HCl and extracted with CH_2_Cl_2_ (3X). The organics were combined, dried over Na_2_SO_4_, and concentrated *in vacuo*. The crude was then purified via silica gel chromatography (0 to 50% EtOAc in Hexanes, 15 CV) to yield (*S*)-N-(1-cyclopropylethyl)-2-methoxynicotinamide (298.4 mg, 1.35 mmol, 52% yield) as a beige solid.

(*S*)-N-(1-cyclopropylethyl)-2-methoxynicotinamide (312.3 mg, 0.1.42 mmol, 1.0 eq) was added to a 20 mL vial along with 4M HCl in Dioxane (11 mL, 0.12 M to (*S*)-N-(1-cyclopropylethyl)-2-methoxynicotinamide). The reaction was brought up to 90 °C and stirred overnight. After the overnight, the solvent was removed, the crude was suspended in MeOH then filtered. The precipitate was then purified via silica gel chromatography (0 to 10% MeOH in CH_2_Cl_2_, 15 CV, dry loaded with celite) to yield **N-[(1*S*)-1-cyclopropylethyl]-2-oxo-1H-pyridine-3-carboxamide** (142 mg, 0.69 mmol, 48% yield) as a crystalline, white solid.

^1^H NMR (400 MHz, DMSO-*d*_6_) δ 12.46 (br, 1H), 9.79 (d, *J* = 8.0 Hz, 1H), 8.31 (dd, *J* = 7.1, 2.3 Hz, 1H), 7.74 – 7.61 (m, 1H), 6.46 (dd, *J* = 7.1, 6.3 Hz, 1H), 3.48 (qd, *J* = 8.0, 6.6 Hz, 1H), 1.19 (d, *J* = 6.6 Hz, 3H), 0.93 (dd, *J* = 8.1, 4.9 Hz, 1H), 0.49 – 0.33 (m, 2H), 0.23 (dd, *J* = 16.7, 3.5 Hz, 2H).

Chemical Formula: C_11_H_14_N_2_O_2_; MW: 206.24.

LC/MS (ESI): m/z = 207.21 [M+H]^+^. Retention time on LC/MS: 1.66 min.

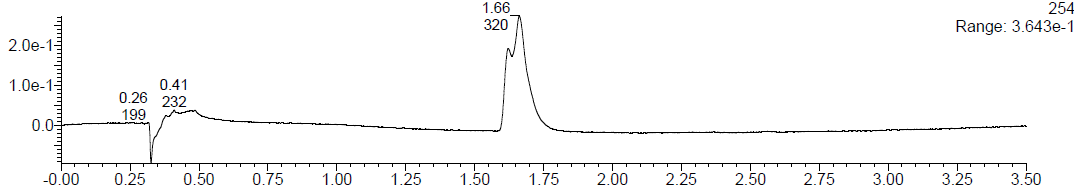

**N-benzyl-2-oxo-1H-pyridine-3-carboxamide**

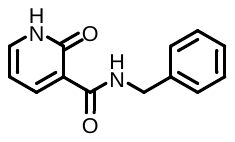

To a 20 mL vial was added 2-methoxynicotinic acid (200 mg, 1.31 mmol, 1.0 eq.) and HATU (521 mg, 1.37 mmol, 1.05 eq). Dry MeCN (2 mL, 0.7 M to 2-methoxynicotinic acid) and NEt_3_ (400 µL, 2.87 mmol, 2.20 eq.) were then added, and the reaction was stirred at room temperature for 15 min. After, phenylmethanamine (143 µL, 1.31 mmol, 1.0 eq) was added to the vial and the reaction was allowed to stir overnight at room temperature. The next morning, the reaction was quenched into 1M HCl and extracted with CH_2_Cl_2_ (3X). The organics were combined, dried over Na_2_SO_4_, and concentrated *in vacuo*. The crude was then purified via silica gel chromatography (0 to 50% EtOAc in Hexanes, 15 CV) to yield N-benzyl-2-methoxynicotinamide (234.7 mg, 0.97 mmol, 74% yield) as a white solid.

N-benzyl-2-methoxynicotinamide (234.7 mg, 0.96 mmol, 1.0 eq) was added to a 20 mL vial along with 4M HCl in Dioxane (8 mL, 0.12 M to N-benzyl-2-methoxynicotinamide). The reaction was brought up to 90 °C and stirred overnight. After the overnight, the solvent was removed, the crude was suspended in MeOH then filtered. The precipitate was washed with more MeOH and then dried to yield **N-benzyl-2-oxo-1H-pyridine-3-carboxamide** (66 mg, 0.29 mmol, 30% yield) as a chalky, white solid.

^1^H NMR (400 MHz, Chloroform-*d*) δ 11.94 (br, 1H), 9.91 (br, 1H), 8.66 (d, *J* = 7.2 Hz, 1H), 7.47 (d, *J* = 6.0 Hz, 1H), 7.40 – 7.28 (m, 5H), 6.53 (t, *J* = 6.8 Hz, 1H), 4.68 (d, *J* = 5.8 Hz, 2H).

Chemical Formula: C_13_H_12_N_2_O_2_; MW: 228.25.

LC/MS (ESI): m/z = 229.22 [M+H]^+^. Retention time on LC/MS: 1.68 min.

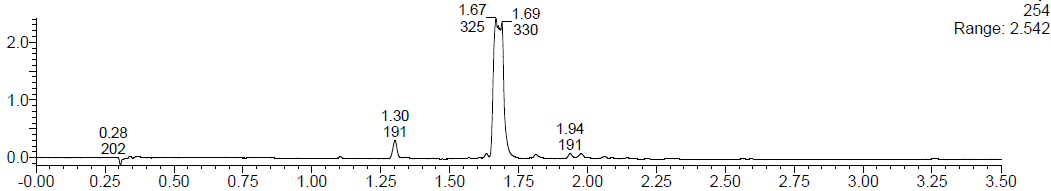

**2-(3-aminophenoxy)-1-pyrrolidin-1-ylethanone**

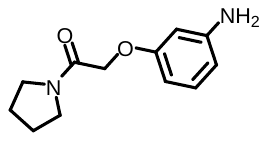

To a 20 mL vial was added 2-(3-nitrophenoxy)acetic acid (3.0 g, 15.0 mmol, 1.0 eq.) and HATU (6.1 g, 16.0 mmol, 1.05 eq). Dry MeCN (11 mL, 1.4 M to 2-(3-nitrophenoxy)acetic acid) and NEt_3_ (4.7 mL, 33.0 mmol, 2.20 eq.) were then added, and the reaction was stirred at room temperature for 15 min. After, pyrrolidine (1.4 mL, 17.0 mmol, 1.1 eq.) was added to the vial and the reaction was allowed to stir overnight at room temperature. The next morning, the reaction was quenched into 1M HCl and extracted with CH_2_Cl_2_ (3X). The organics were combined, dried over Na_2_SO_4_, and concentrated *in vacuo* onto celite. The crude was then purified via silica gel chromatography (0 to 10% MeOH in CH_2_Cl_2_, 15 CV, dry loaded with celite) to yield 2-(3-nitrophenoxy)-1-(pyrrolidin-1-yl)ethan-1-one (3.6 g, 14.0 mmol, 95% yield) as a brown solid.

2-(3-nitrophenoxy)-1-(pyrrolidin-1-yl)ethan-1-one (2.3 g, 9.2 mmol, 1.0 eq.) was added to a 50 mL flask along with Tin(II) Chloride dihydrate (6.2 g, 28.0 mmol, 3.0 eq.). Add dry EtOH (10 mL, ~1.0 M to 2-(3-nitrophenoxy)-1-(pyrrolidin-1-yl)ethan-1-one)) and then reflux overnight. After overnight, filter to remove any tin solids. Then dump filtrate into a separatory funnel with a large amount of brine, extract with DCM (3X), and then dry the combined organics over Na_2_SO_4_, and concentrated *in vacuo* onto celite. The crude was then purified via silica gel chromatography (0 to 10% MeOH in CH_2_Cl_2_, 25 CV) to yield **2-(3-aminophenoxy)-1-pyrrolidin-1-ylethanone** (349.3 mg, 1.58 mmol, 17% yield) as a brown foamy solid.

^1^H NMR (400 MHz, Chloroform-*d*) δ 7.05 (t, *J* = 8.3 Hz, 1H), 6.37 – 6.30 (m, 3H), 4.58 (s, 2H), 3.55 – 3.49 (m, 4H), 1.94 (q, *J* = 6.8, 6.1 Hz, 2H), 1.88 – 1.80 (m, 2H).

Chemical Formula: C_12_H_16_N_2_O_2_; MW: 220.27.

LC/MS (ESI): m/z = 221.26 [M+H]^+^. Retention time on LC/MS: 1.11 min.

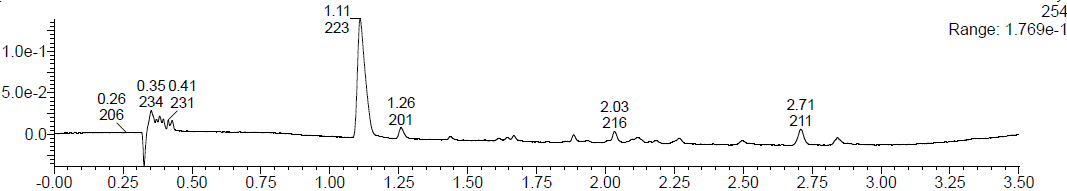

^1^H NMR of **5-(2-methoxyphenyl)-1,3,4-oxadiazol-2-amine**
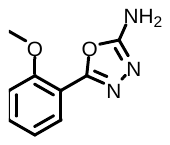

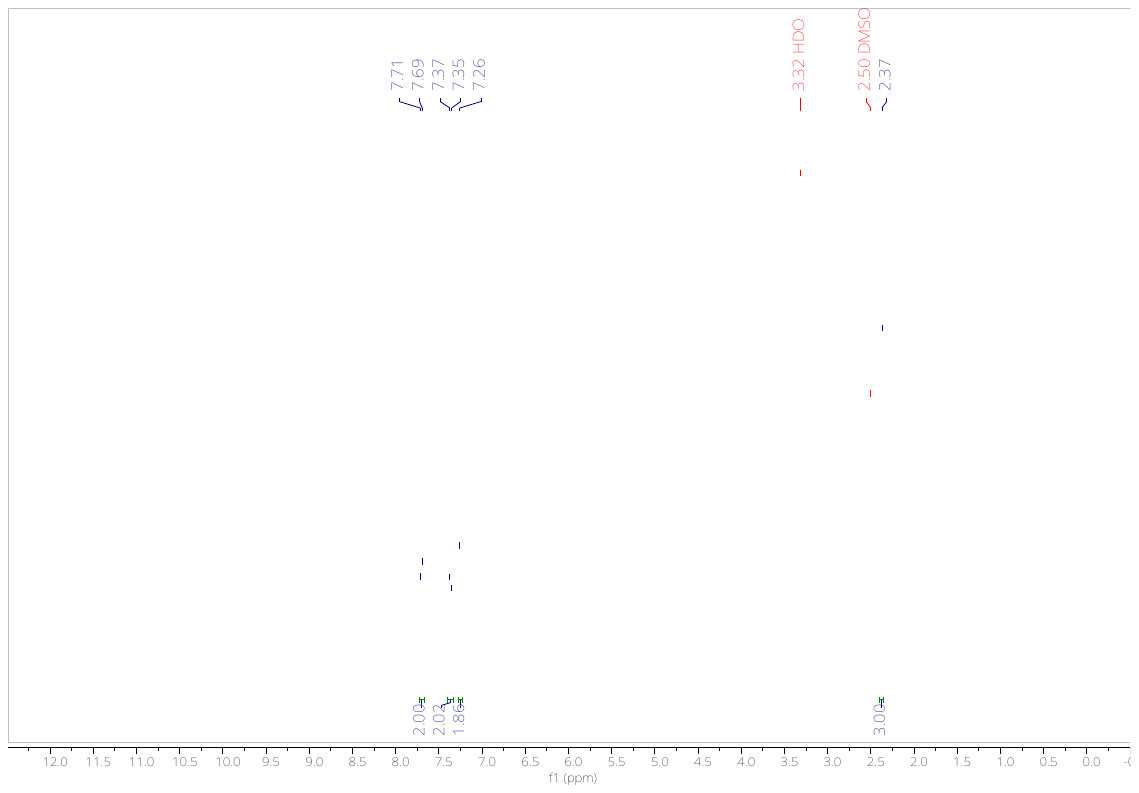

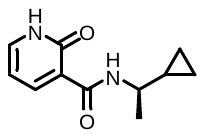
^1^H NMR of **N-[(1R)-1-cyclopropylethyl]-2-oxo-1H-pyridine-3-carboxamide**
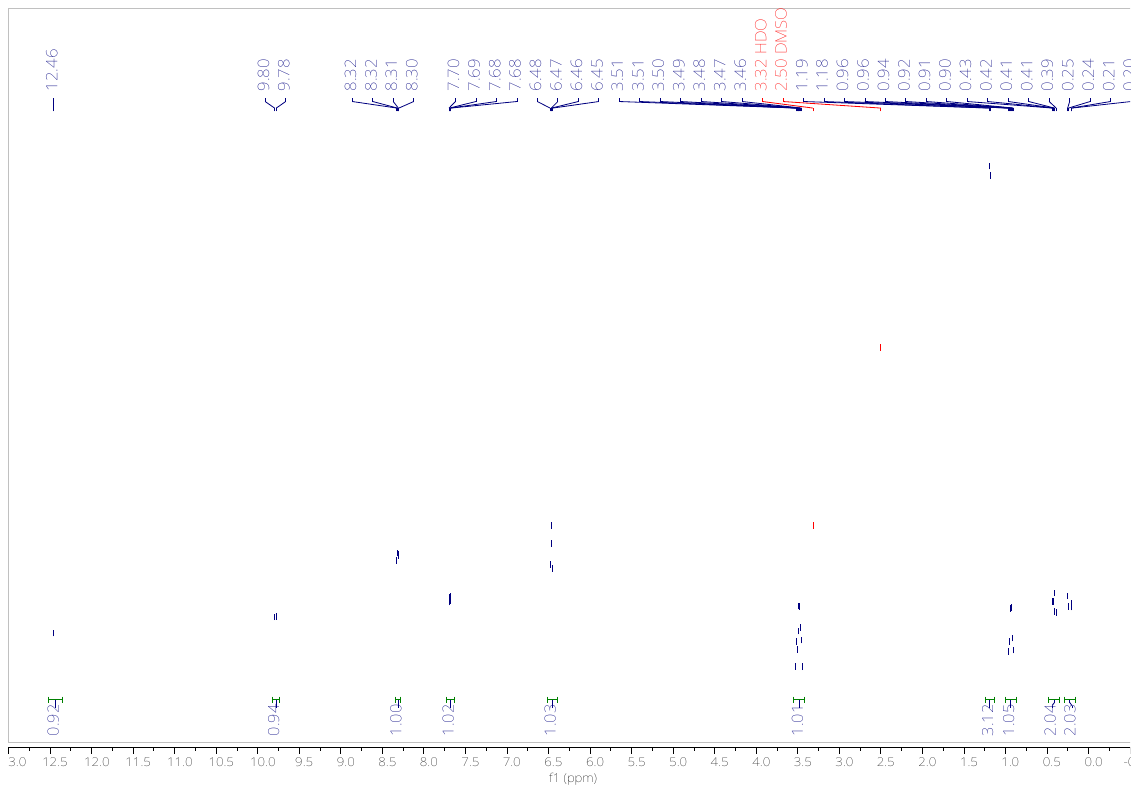

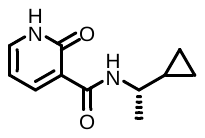
^1^H NMR of **N-[(1S)-1-cyclopropylethyl]-2-oxo-1H-pyridine-3-carboxamide**
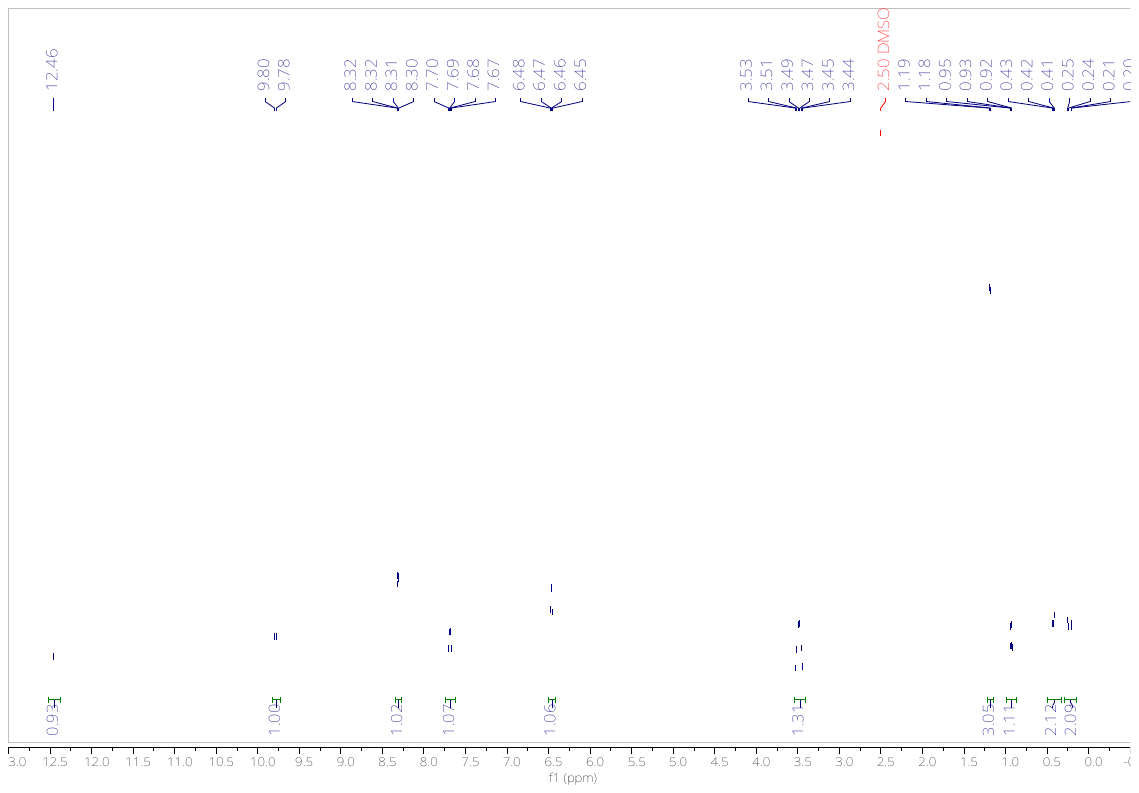

^1^H NMR of **N-benzyl-2-oxo-1H-pyridine-3-carboxamide**

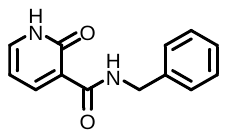

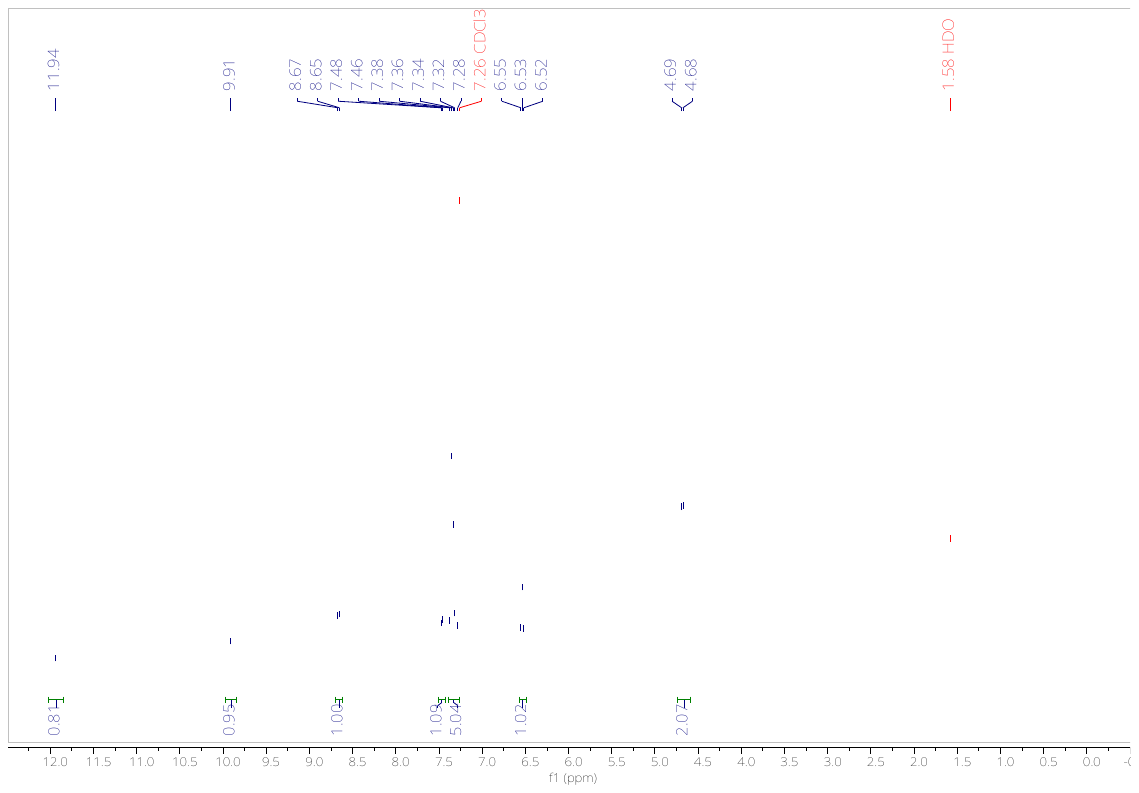

^1^H NMR of **2-(3-aminophenoxy)-1-pyrrolidin-1-ylethanone**

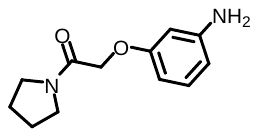

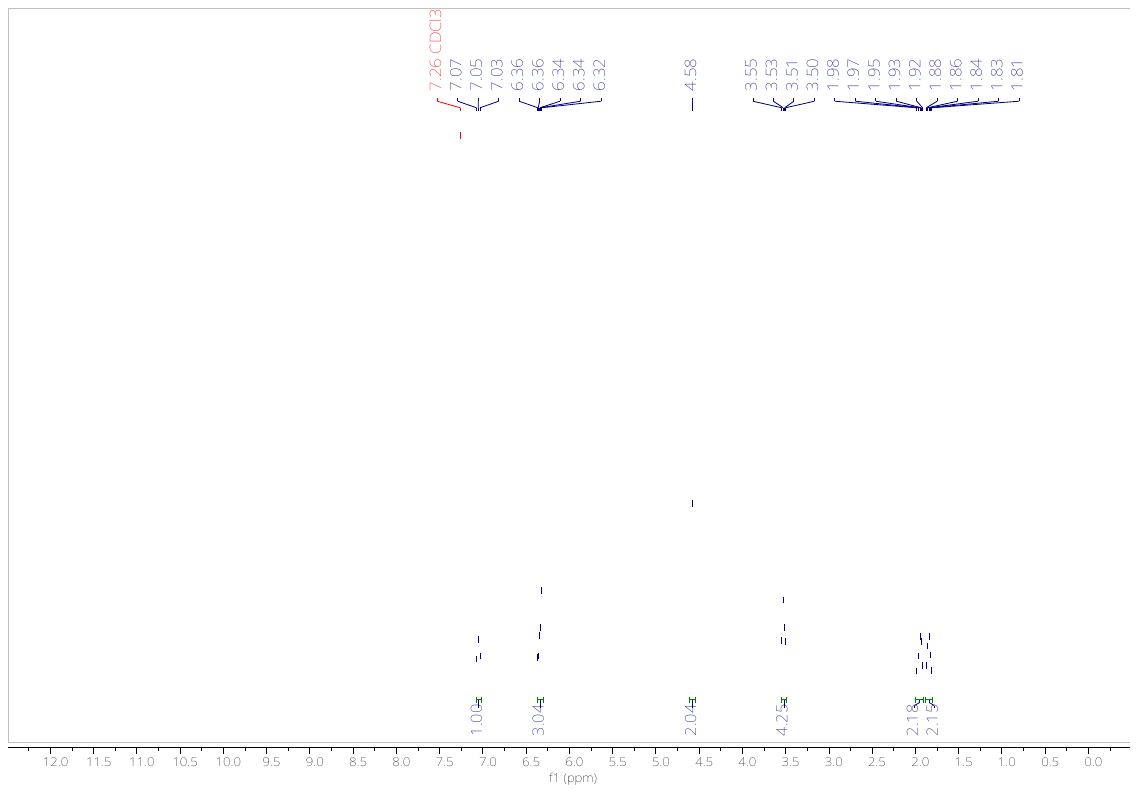
